# Early embryonic heat shock induces long-term epigenetic memory by affecting the transition to zygotic independence

**DOI:** 10.1101/2021.10.27.466124

**Authors:** Lovisa Örkenby, Signe Skog, Helen Ekman, Unn Kugelberg, Rashmi Ramesh, Marie Roth, Daniel Nätt, Anita Öst

## Abstract

Early-life stress can generate persistent life-long effects that impact adult health and disease risk, but little is known of how such programming is established and maintained. Previous use of the *Drosophila* strain *w^m4h^* show that an early embryonic heat shock result in stable epigenetic alteration in the adult fly. To investigate the potential role of small non-coding RNA (sncRNA) in the initiation of such long-term epigenetic effects, we here generated a fine timeline of sncRNA expression during the first 5 stages of *Drosophila* embryogenesis in this strain. Building on this, we show that (1) miRNA is increased following early embryonic heat shock, and (2) the increased miRNA is coming from two separate sources, maternal and zygotic. By performing long RNA sequencing on the same single embryo, we found that a subgroup of miRNA with maternal origin, had a strong negative correlation with a group of early zygotic transcripts. Critically, we found evidence that one such early zygotic transcript, the insulator binding factor Elba1, is a Su(var) for *w^m4h^*. The findings provide insights of the dynamics and stress-sensitivity of sncRNA during the first embryonic stages in *Drosophila* and suggest an interplay between miRNA, Elba1 and long-term epigenetic alteration.

**HIGHLIGHTS:**

- We provide a high-resolution timeline for sncRNA for *Drosophila* stage 1-5 embryos
- Heat shock before midblastula transition (MBT) results in a massive upregulation of miRNA at cellularization
- Heat shock-induced miRNAs negatively associate with downregulation of a specific subset of pre-MBT genes
- Elba1 is a position-effect-variegation (PEV) modifier for *w^m4h^*
- Heat shock-induces an “leaky” expression of genes that overlap with Elba 1-3 binding sites

## INTRODUCTION

Early-life is carefully orchestrated by a plethora of processes that allows for both developmental robustness and plasticity, ultimately regulating the diversity of phenotypes from a single genome. This provides the foundation for the Developmental Origins of Health and Disease (DOHaD) hypothesis, which postulates that the etiologies of major public health issues, such as obesity, type 2 diabetes and heart disease, depend on sub-optimal conditions during sensitive periods early in life (Suzuki, 2018). In humans, this can be caused by factors such as malnutrition, smoking, physical or psychological trauma (reviewed in (Block and El-Osta, 2017; Cunliffe, 2016; Knopik et al., 2012; Wong and Langley, 2016)), in *Arabidopsis* by hyperosmotic stress (Sani et al., 2013) and in *Drosophila melanogaster* by e.g. heat shock (Seong et al., 2011). The developmental timing of exposure has proven crucial for determining the outcome and, to date, it is poorly understood what sets such sensitive developmental periods apart from insensitive. Moreover, the molecular mechanisms initiating and shaping the response, as well as how memories of these exposures are kept throughout the developmental reorganization of the chromatin landscape, remains to be understood.

In *Drosophila*, embryogenesis is extremely rapid with less than 3 h from fertilization to gastrulation and only 24 h to the first larvae stage. The main reason for this is that the early *Drosophila* embryo, like most insects, undergoes a series of rapid mitotic events without cytokinesis where all nuclei share the same cytoplasm (reviewed in (Hamm and Harrison, 2018)). These cycles, which are only separated by a few minutes, are too short for extensive zygotic transcription (De Renzis et al., 2007; Kwasnieski et al., 2019) making these pre-cellular stages of *Drosophila* embryogenesis highly dependent on maternally loaded proteins and RNAs. At the midblastula transition (MBT), there is a lengthening and synchronization of mitotic cycles that coincides with the zygotes claim of transcriptional independence, a process crucial for the maternal to zygotic transition (MZT) (Vastenhouw et al., 2019). Before MZT, there are no higher order of chromatin organization reported. During MZT however, several well-coordinated events, driven by an interplay between maternally provided products and zygotic *de novo* transcription, lead to establishment of chromatin states and a chromosomal 3D architecture that can be detected by Hi-C as topologically associated domains (TADs) (Hamm and Harrison, 2018; Hug et al., 2017; Li et al., 2014; Stadler et al., 2017; Yuan et al., 2016). One important component for the establishment of higher order chromatin structure are insulator binding factors that binds to genomic cis-regulatory insulator sequences to prevent leakage of the regulatory environment between neighboring genes and across longer distances (Stadler et al., 2017). Recently, a family of insulator binding proteins was discovered, the Elba complex, that is expressed just before the MBT to ensure partition of transcription units during the transition to zygotic independence (Aoki et al., 2012; Ueberschär et al., 2019).

The role of microRNA (miRNA) as one of the zygotic key regulators for the *Drosophila* MZT by clearing maternal transcripts has been known for more than a decade (Bushati et al., 2008). In addition, miRNA together with other small non-coding RNAs (sncRNAs), including piRNA, fragments of tRNA (tsRNA), and rRNA (rsRNA), play important roles in inter- and transgenerational epigenetic inheritance (de Castro Barbosa et al., 2015; Grandjean et al., 2015; Nätt et al., 2019; Sharma et al., 2016; Zhang et al., 2018). In combination with the known involvement of siRNA in heterochromatin formation (Li et al., 2009), piRNA in transposon silencing (Huang et al., 2013), and certain tRNA halves in regulating histone biogenesis (Boskovic et al., 2019), it is easy to envision a general role for sncRNA, including miRNA, in initiating or influencing the early higher order chromatin landscape (Allshire and Madhani, 2018; Holoch and Moazed, 2015; Johnson and Straight, 2017). Furthermore, cellular responses to stress involves upregulation and activation of specific miRNA (Leung and Sharp, 2010; Olejniczak et al., 2018) and proteins (Chen et al., 2018), as well as fragmentation of tRNA (Thompson et al., 2008). Thus, in addition to a central role in initiating chromatin states, sncRNA plays a vital role in the cellular stress response. Currently, there are no fine-resolution data of sncRNA covering the first stages of embryogenesis.

Here, we explore the effects of environmental stress on the expression of sncRNA during early *Drosophila* embryogenesis. We specifically aimed to identify sensitive developmental windows in which stress might induce long-lasting memories. Furthermore, by examining gene- and sncRNA expression within the same single *Drosophila* embryos, we aimed to identify critical interactions between sncRNA and gene regulatory networks in such a sensitive window.

As previously shown (Bughio et al., 2019; Hartmann-Goldstein, 1967; Lu et al., 1998; Seong et al., 2011), we find that heat shock before the MBT reduces the epigenetic-mediated, H3K9/H3K20 methylation-dependent silencing of the position effect variegation (PEV) *white* gene, which is an adult eye-color heterochromatin reporter (Elgin and Reuter, 2013). Early heat shock further results in retention of maternally loaded miRNA’s, including a specific group of miRNA that negatively associates with the expression of a specific group of early zygotic transcripts. Finally, frame-shift mutation of one of these genes, *Elba1* (a.k.a. *Bsg25A*), efficiently mimicked the effect of heat shock on the adult eye color reporter, thus suggesting that a temporal expression of embryonic insulators have long-lasting epigenetic effect.

## RESULTS

### Heat shock during the first two hours of embryogenesis causes long-term effects

To identify sensitive periods during *Drosophila* development, where a stress exposure can induce a long-term memory, we used the position-effect-variegation strain *w^m4h^* (Figure 1A). This strain has an inversion on the X chromosome, positioning the *white* gene close to the pericentric heterochromatin. The expression of *white*, which is needed for eye pigmentation, is therefore controlled by the centromeric chromatin state and the adult eye color can be used as a reporter for heterochromatin at this locus (Elgin and Reuter, 2013). Variegation of *w^m4h^* is controlled by the methyltransferases Su(var) 3-9, Su(var) 4-20, Su(var)2-5 and E(z) (Phalke et al., 2009), and we previously showed that this reporter is sensitive to paternal diets (Öst et al., 2014).

**Figure 1.**
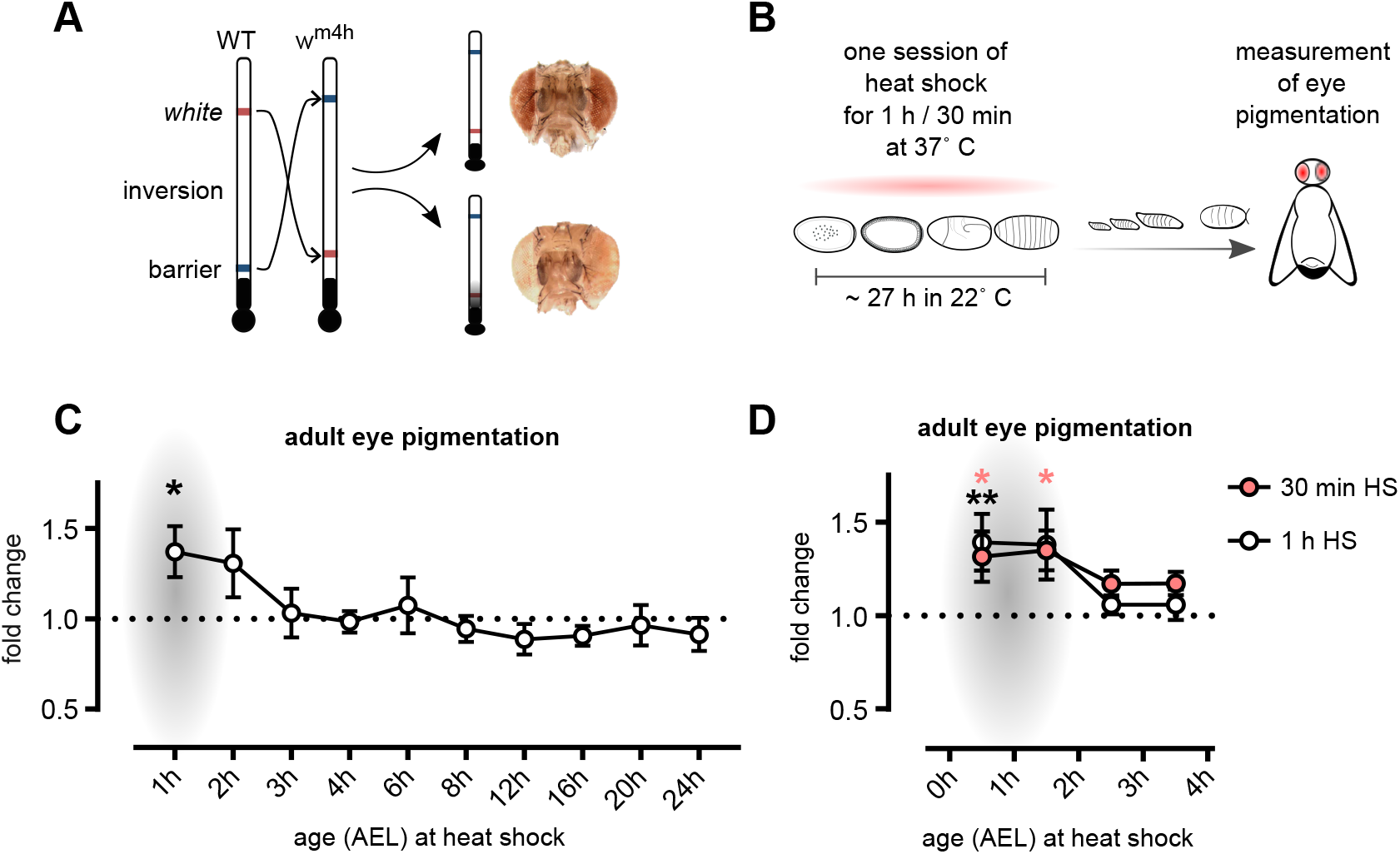
The most sensitive period for heat shock-induced epigenetic programming is the first two hours of embryogenesis. **A.** *Drosophila w^m4h^* has an inversion of the *white* gene, needed for eye pigmentation, which places this gene in proximity to the centromeric heterochromatin. This enables detection of heterochromatin spreading through measurement of eye pigmentation. **B.** Eggs were collected in 1 h intervals and exposed to one session of 1 or 0.5 h heat shock at different developmental time windows, or kept as controls. Developing flies were kept in 22°C until pupae hatching and eye pigmentation was measured in 5 days old males. **C-D.** Eye pigmentation in relation to controls (not exposed to heat shock). The most stress-sensitive period was found during the first hour of embryogenesis. At this time-point, a 1 h (C-D) or 30 min (D) heat shock resulted in adult flies with more pigmented eyes, indicative of a more open chromatin at this locus. Heads were measured in groups of 3-10 and normalized to the average optical density per head. n ≥ 7 (C) and n ≥ 5 (D) pools per time window. AEL = after egg laying. Presented with ± SEM, * (adjusted p ≤ 0.05) and ** (adjusted p ≤ 0.01) with ordinary one-way ANOVA with Dunnett’s multiple comparison test.

To enable a high-resolution mapping of sensitive periods for environmentally induced memories, we used heat shock. In contrast to other environmental challenges such as suboptimal nutrition and exposure to toxins, heat shock allows for a sharp and distinct intervention time. We performed a one-hour heat shock during different time points of *Drosophila* development, throughout embryogenesis (Figure 1B-C), as well as during the larva stages (Figure S1). As *white* expression peaks during pupation, we assessed eye color in 5 day old male adults. We found that the first two hours in embryogenesis are the most sensitive period for heat shock induction of long-term effects on heterochromatin, as assayed by *w^m4h^* variegation (Figure 1C, S1), and confirmed this sensitive time period using an even shorter heat shock exposure time (30 min) (Figure 1D). Our finding is consistent with previous work identifying the first 0-3 hours after fertilization as a time in which the epigenome is sensitive to heat shock stress (Bughio et al., 2019; Hartmann-Goldstein, 1967; Lu et al., 1998; Seong et al., 2011). Importantly, we performed heat shock in more narrow intervals, and while we did get effects in 0-1 h, and 1-2 h, we did not detect any significant long-lasting effects on *white* expression in 2-3 h old embryos, indicating that in order for long-term effects to occur, the exposure of a stressor must happen before the MBT.

### Early *Drosophila* embryogenesis is accompanied by dynamic shifts in sncRNA

The first two hours of *Drosophila* embryogenesis, entailing stage 1-3 (at 22° C), is characterized by rapid mitotic cycles dependent on maternally loaded mRNAs and proteins (Figure 2A) (Bushati et al., 2008; Tadros and Lipshitz, 2009; Vastenhouw et al., 2019). In concordance with the rapid cell divisions, there is no clear higher order chromatin architecture in this period (Hug et al., 2017). Chromatin states are established at MBT, around stage 5, when the tempo of mitosis subsides and zygotic transcription is activated (Rudolph et al., 2007). As sncRNAs are highly present already in stage 1-3 and are known to modulate higher order chromatin structure, we hypothesized that heat shock-induced changes of sncRNA might precede and guide later *de novo* heterochromatin formation. As there are no high-resolution time courses of changes in sncRNA during these early stages of *Drosophila* embryogenesis, we began by performing sncRNA sequencing on single embryos, embracing stage 1-5 (Figure 2A). Bioinformatic analysis showed that the relative proportions (Figure 2B) and size distributions (Figure 2C) of different sncRNA classes were highly dynamic during stages 1-5. Thus, early embryogenesis is accompanied by rapid temporal changes in the sncRNA-profile.

**Figure 2.**
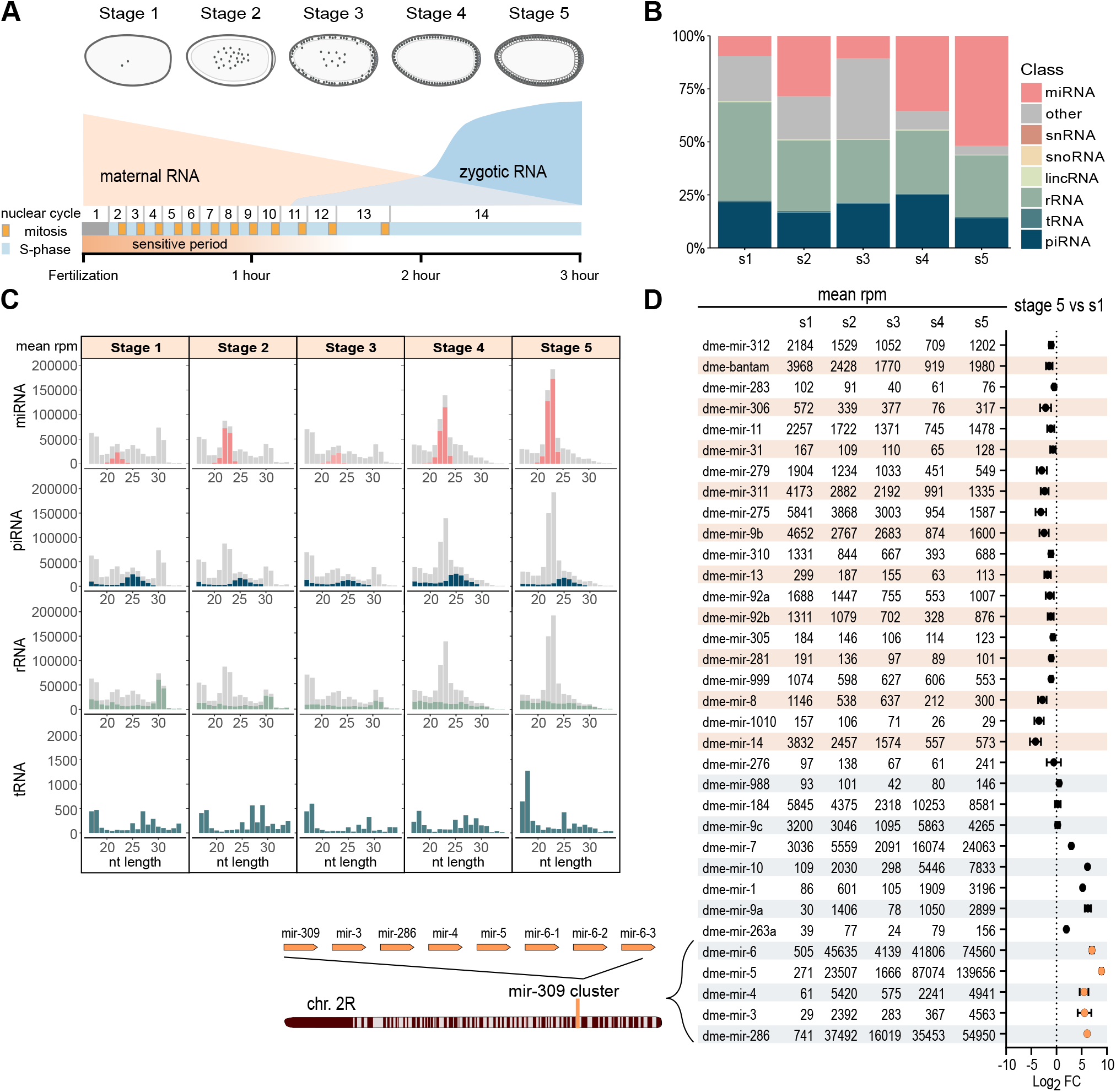
Early embryogenesis is characterized by rapid changes of sncRNA. **A.** Schematic illustration. The early *Drosophila* embryo undergoes a series of rapid mitotic events without cytokinesis where the pre-cellular stages of *Drosophila* embryogenesis are dependent on maternally loaded proteins and RNAs. Zygotic gene activation occurs in two waves, one minor during stage 3 at pole cell formation and one after cellularization in stage 5. **B.** Relative proportions of sncRNA classes in *w^m4h^* embryos during stage 1-5. **C.** Mean rpm per nucleotide length and stage. **D.** (left) miRNA expression (mean rpm) per stage, beige = maternal and light blue = zygotic. (right) Log2 fold change of rpm per indicated miRNA between stage 5 embryos and the mean rpm of stage 1 embryos. (bottom left) Schematic illustration of mir-309 cluster. n = 5 of stage 1-3 and n = 4 of stage 4 and 5.

As expected during the maternal to zygotic transition, stage 3 is characterized by an increase of degradation products (“other”, Figure 2B) which then decrease at the end of MZT. There are also more rsRNAs in stage 1 than in the other stages, while tRNA fragments (tsRNAs) are expressed at low levels at all stages (Figure 2B-C). In contrast, piRNAs are highly expressed throughout all stages. The most striking change, however, was in miRNA levels during early embryogenesis, with a distinct peak in stage 2 followed by a decline in stage 3 and then a rise again in stages 4 and 5.

While multiple miRNAs were reduced during this developmental time window, suggestive of a maternal origin, the zygotic miR-309 cluster showed a pronounced upregulation (Figure 2D, S2, S3). Controlled by Zelda, a maternally provided pioneering transcription factor (Fu et al., 2014; Liang et al., 2008), the miR-309 cluster plays an important role in the zygotic driven pathway that degrades maternal transcripts (Bushati et al., 2008). Sequential comparison of stage 2 against 1, 3 against 2, 4 against 3, and 5 against 4 revealed that even though transcripts from the miR-309 cluster had a sharp increase in stage 4 and 5 (Figure S2C-D), the most significant change was between stage 1 and 2 (Figure S2A-E). To test that the early increased expression of this cluster was not an artifact of a few outlier miRNA sequences, we compared unique miRNA sequences per sample and stage (Figure S3). This revealed a uniform upregulation of the miR-309 cluster and strengthened the notion that there in fact is an upregulation of this cluster between stage 1 and 2. Previous findings have shown that members of this cluster are expressed at low levels in 0-1 hour old embryos and are strongly induced 2-3 hours after egg laying (Aravin et al., 2003; Bushati et al., 2008; Fu et al., 2014; Ninova et al., 2014; Ruby et al., 2007). Furthermore, low levels of Zelda has been detected in the nucleus already at nuclear cycle 2 (Nien et al., 2011). Our results align with these findings and support a scenario where the miR-309 cluster is starting to be transcribed in low levels already between embryonic stage 1 and 2.

### Heat shock in the sensitive period results in a rapid change of sncRNA at the time of heterochromatin formation

To investigate if a heat shock in our sensitive period results in changes to the sncRNA-profile at cellularization and heterochromatin formation, we next heat shocked 0-0.5 hour old embryos for 30 min and then aged them to stage 5 before collecting single embryos for sequencing (Figure 3A). As we noticed before (Figure 2B-C), the sncRNA profiles of stage 5 embryos are dominated by miRNAs with the expected length of 22-24 nucleotides (Figure 3B-C). We did not detect any changes in size distribution between conditions, but we found a significant increase of miRNAs in heat shocked samples (Figure 3D). Correlation plots of different sncRNA classes between heat shocked and control embryos confirmed that a high extent of unique miRNA sequences were significantly upregulated (Figure 3E-F). In addition, correlation plots and differential expression analysis revealed a similar upregulation of piRNA (Figure 3E-F), while rsRNA and tsRNA showed diverse responses (Figure 3E-F). As expected, we found that heat shock induced more tRNA halves originating from the 5’ terminal of mature Gly-GCC (Figure S4, top). This tsRNA, which is also called tRNA-derived stress-induced small RNA, is cleaved at the anti-codon loop by ribonucleases like angiogenin, and is generated in response to different kinds of cellular stress leading to the formation of stress granules (Emara et al., 2010). In sperm, such tRNA 5’ halves are also affected in response to paternal diet resulting in metabolic changes in the offspring (Chen et al., 2016; Nätt and Öst, 2020), but their role in early embryogenesis in *Drosophila* is still unknown. Nonetheless, the most prominent effect of early embryonic heat shock on stage 5 embryos was the significant increase of 184 unique miRNA sequences, indicative a miRNA dependent stress response proceeding MBT and zygotic gene activation (ZGA).

**Figure 3.**
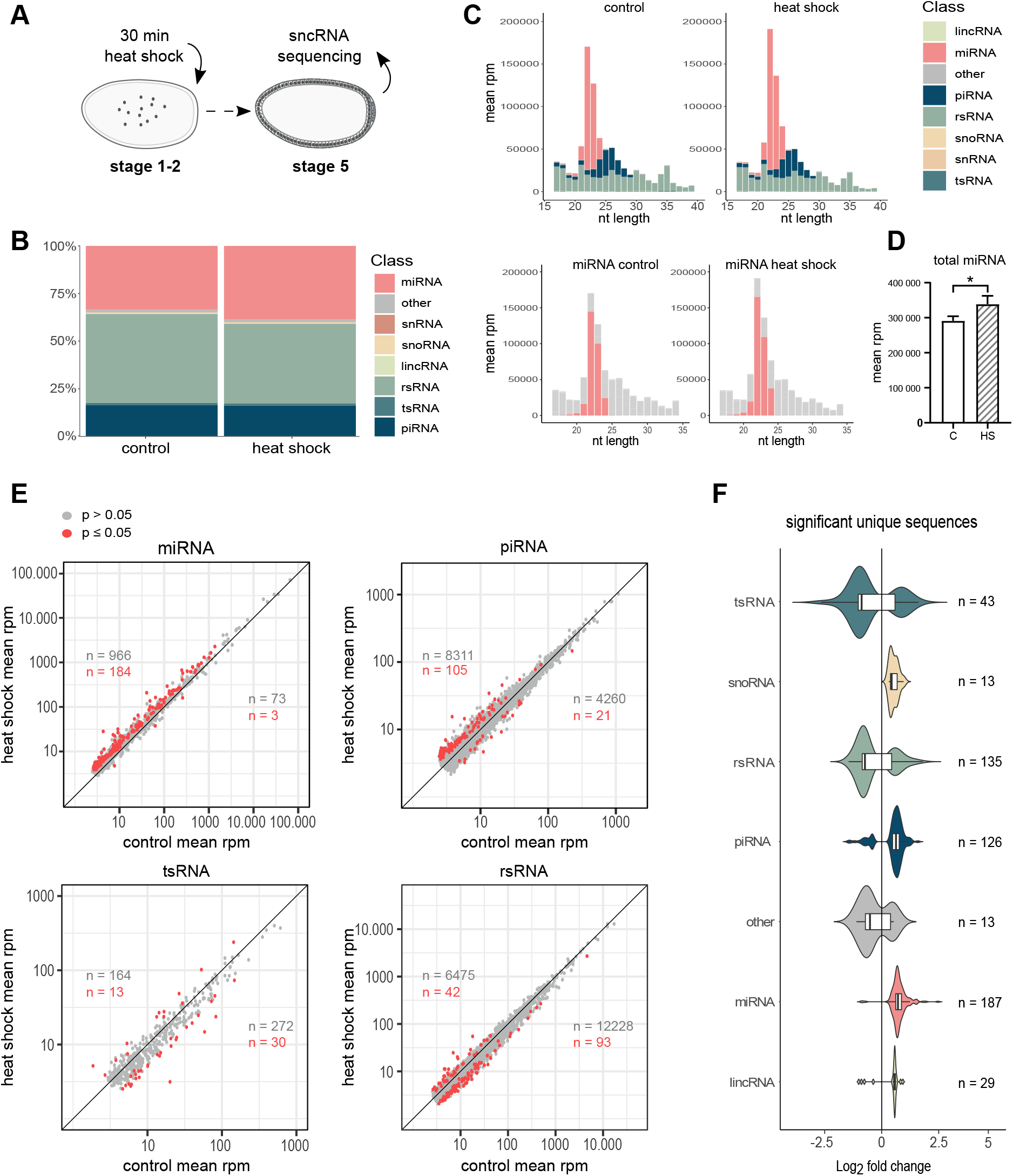
Heat shock in sensitive period leads to upregulation of miRNA. **A.** *w^m4h^* eggs were collected in short intervals (30 min) and immediately heat shocked for 30 min at 37° C (or kept as controls). All eggs were aged for approximately two hours in 22° C, dechorionated and manually staged under microscope and collected at stage 5. **B.** Relative proportions of each sncRNA class within the two conditions. A greater proportion of miRNA and a smaller proportion of rsRNA was found in embryos exposed to heat shock during the sensitive period. **C.** Read-length distribution per treatment group and sncRNA class (top), or only miRNA (bottom, red). The majority of increased miRNAs in heat shocked samples came from 22-24 nt reads. **D.** Expression of total miRNA between conditions. The miRNA levels are significantly increased in heat shocked samples. Presented with ± SEM, * = p < 0.05 using unpaired one-tailed t-test. **E.** Scatter plots comparing rpm normalized expression levels of unique miRNA, piRNA, tRNA- or rRNA fragments between control and heat shocked embryos. Red = FDR corrected p ≤ 0.05, grey = FDR corrected p > 0.05. **F.** Log2 fold change of significantly (FDR corrected p ≤ 0.05) expressed unique reads per sncRNA class between heat shocked and control samples. 24 single embryos were sequenced per condition.

The general upregulation of miRNA could be due to either increased zygotic transcription or decreased degradation of maternal transcripts. These two scenarios were discriminated by comparing the miRNA profiles from stage 1-5 embryos (Figure 2D, S2-3). Since we found a clear distinction of miRNA species found at the two first stages compared to stage 4-5 (Figure 2D, S3), we classified miRNAs having their peak expression during stage 1-2 as maternally loaded and miRNAs that had their peak in later stages (or peak during the minor wave of ZGA) to be zygotic. Using this division, we found that the upregulation of miRNA in response to heat shock arises from the combination of both increased zygotic transcription (Figure 4A, C) and loss of degradation of maternally provided miRNAs (Figure 4A-B). Importantly, we did not detect any difference in the expression of the mir-309 cluster, accounting for 69-75 % of total miRNA at stage 5 (Figure 4D). This suggests that heat shock targets the upregulation of a specific group of zygotic miRNAs.

**Figure 4.**
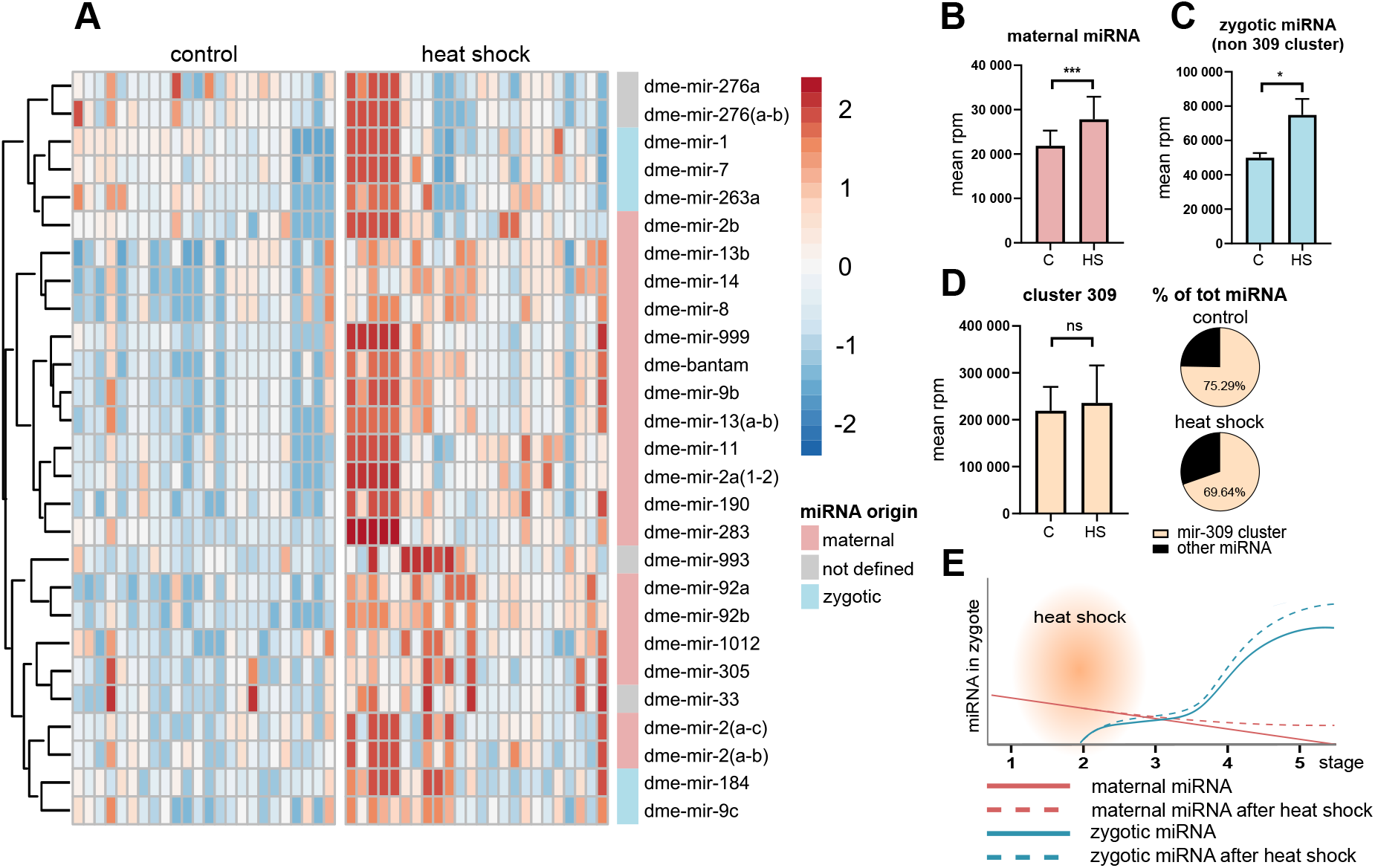
Heat shock during sensitive period leads to abnormal abundance of maternal miRNA in stage 5. **A.** Heatmap of significantly (FDR corrected p ≤ 0.05) expressed miRNA between heat shocked and control embryos, 24 embryos per condition. Color bar represents the relative rpm expression per miRNA. **B-C.** Mean expression of maternal (B) and zygotic (C) miRNAs, where reads aligning with the mir-309 cluster were excluded. **D.** Mean expression of reads mapping to the mir-309 cluster. **E.** Schematic illustration of maternal and zygotic miRNA levels during the first 5 stages of embryogenesis with or without heat shock exposure. The illustration is based on our observations from miRNA expression profiles during stage 1-5 (Figure 2) and B-C. In bar charts: * = p ≤ 0.05, **** = p ≤ 0.0001, ns = non-significant, with unpaired two-tailed t-test. Error bars represents SD and 24 embryos were analyzed per condition.

In all, an early embryonic heat shock results in retention of maternally loaded miRNAs as well as specific upregulation of zygotic miRNAs that together contributes to a pronounced increase of miRNA before gastrulation (Figure 4E).

### Heat shock induced changes in miRNA correlates with the downregulation of selected pre-MBT genes

Given that the chromatin architecture can be influenced during the embryonic pre-ZGA period, we next investigated if excessive miRNA during this period correlated with changes to epigenetic factors controlling chromatin states. Since the *w^m4h^* locus is known to be controlled by classical H3K9me3-dependent mechanisms, we hypothesized that the upregulation of miRNA would result in downregulation of factors involved in this mechanism. To test this hypothesis, we prepared libraries for ribo-minus RNA-seq using the same single-embryo RNA-extractions that were used for our sncRNA analysis (Figure 5A).

**Figure 5.**
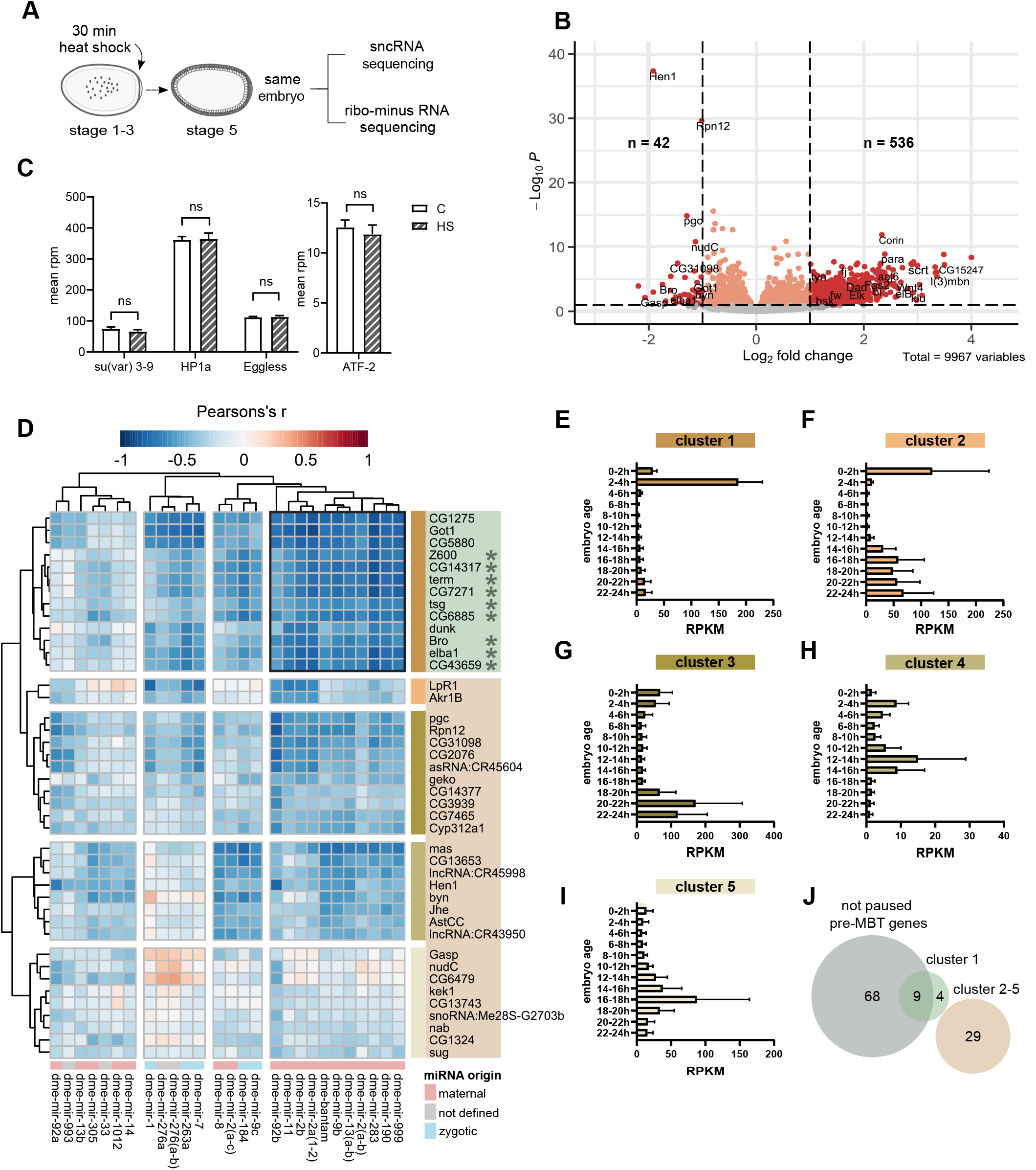
Heat shock reduces specific embryonic pre-MBT genes. **A.** Ribo-minus libraries for long RNA sequencing were prepared from the same RNA extracts as used for sncRNA sequencing of heat shocked and control embryos. **B.** Volcano plot showing differentially expressed long RNA. Dark red indicates significance at p ≤ 0.05 (FDR corrected p-values) and a log2 fold change ≥ ± 1. **C.** No significant changes of Su(var) 3-9, Eggless, HP1a or ATF-2 was detected after early embryonic heat shock. **D.** Pearson’s r for all significant upregulated miRNA and downregulated long RNA. Unsupervised Euclidean clustering shows that several maternal miRNAs correlates strongly negative to gene cluster 1. **E-I.** Temporal expression of each gene cluster using modENCODEs data (D). Cluster 1 is specifically expressed at 2-4 h after egg laying, corresponding with the pre-midblastula transition (pre-MBT) and midblastula transition (MBT) (E). Cluster 2 and 3 has their peak expression in young embryos followed by a quick decline and re-expression during late embryogenesis (F-G). Genes in cluster 4 shows an unspecific trend (H), whilst cluster 5 peaks in 16-18 h old embryos. **J.** Overlap between gene clusters and staged embryonic data from Chen et al. (2013) shows that cluster 1 mostly consists of not pol II paused pre-MBT genes.

Differential expression analysis showed that heat shock significantly altered the expression of several mRNAs (figure 5B), primarily resulting in gene upregulation (1134 up vs 459 down). We could, however, not detect any significant changes in the expression of H3K9me3-related epigenetic enzymes such as Su(var)3-9, HP1, Egg or ATF-2 (figure 5C). Instead, functional analysis showed that heat shocked embryos had an elevated expression of genes involved in tissue differentiation (Figure S5A), whereas no associations with biological processes were observed in the down- regulated genes. Closer inspection revealed, however, that two of these less expressed genes—early boundary activity 1 (Elba1 also known as Bsg25A) and polar granule component (pgc)—are involved in chromatin organization (Figure S5B).

Unsupervised correlation and clustering analysis between the upregulated miRNA and downregulated long RNA revealed a cluster with strong inverse correlation (Figure 5D, cluster 1; S6 and Supplemental Table 6). This cluster consisted of 11 miRNA, including mir-190, mir-2 and bantam, and 13 mRNA. Using modENCODEs temporal expression data for all genes in each RNA cluster (Figure 5E-I) we found that our identified gene cluster is exclusively expressed in 2-4 h old embryos (Figure 5E), which correlates with the pre-midblastula transition (pre-MBT) and midblastula transition (MBT). To get further insight about this cluster, we compared it with carefully categorized pre-MBT genes from Chen et al. (2013) and found that 9 out of 13 genes in this cluster overlapped with non-paused pre-midblastula transition (pre-MBT) genes (Figure 5J). These pre-MBT genes have been shown to have no detected pausing of Pol II and a peak expression at cellularization, just before the MBT. Moreover, they have been shown to have a strong overlap with a small group of genes identified as the very first zygotic transcripts (De Renzis et al., 2007).

To exclude the possibility that the downregulation of a subgroup of the pre-MBT were caused by a mismatch between morphological and transcriptional age of the heat shocked embryos, leading to a technical bias when sampling, we compared the expression of maternally provided and early zygotic genes (transcribed within 1-2 h of embryogenesis). We could not detect any difference of the distribution of either (Figure S7A-B). As the gene expression is very dynamic during embryonic cellularization and ZGA, we further compared gene expression from control and heat shocked embryos to previously published gene expression of nuclear cycle 14 that were chronologically divided into four parts (Lott et al., 2011). By looking at the linear regression of nuclear cycle 14 gene expression, we classified genes as early (slope ≤ −1), late (slope ≥ 1) or stable expressed (slope between −1 and 1) (Figure S7C-D). We could not detect any temporal bias between our groups using this classification (Figure S7E). Further, the stable expression of miRNA from the mir-309 cluster across samples (Figure 4D) and the elevated occurrence of maternally provided miRNAs (Figure 4B) adds to the conclusion that the differences in gene expression following heat shock were not due to sampling error. Thus, we conclude that there is a heat shock-induced upregulation of miRNA that correlates negatively with a specific group of pre-MBT genes suggestive of a stress-miRNA-pre-MBT gene axis.

### The insulator binding complex Elba associates with de-repressed genes and acts as a Su(var) for *w^m4h^*

Next, we focused on genes that had an elevated expression following heat shock (Figure 5B). These genes were predominantly involved in tissue differentiation and morphological processes (Figure S5A). Developmental patterning has previously been shown to be regulated by the Elba complex, and mutation of any member of this complex results in de-repression of genes (Ueberschär et al., 2019). Importantly, Elba1—one of the chromatin-binding components of this complex— was one of the 9 downregulated non-paused pre-MBT genes following heat shock with the highest inverse expression to the upregulated miRNAs (Figure 5D, S6). Thus, we hypothesized that heat shocked embryos may suffer from lower insulating capacity.

Looking more closely at insulator binding factors with different temporal expression profiles during embryogenesis (Figure S8), we found that two of three Elba factors, Elba1 and Elba2, but also two other insulator binding factors, Insv and CP190, were less expressed after heat shock (Figure 6A). Unsupervised cluster analysis of Pearson’s r scores for insulator binding factors and heat shock induced genes revealed four gene clusters of which cluster 2, the largest cluster, had a strong negative correlation to Elba1, Elba2, Insv and CP190 (Pearson’s r: Elba1 mean = - 0.66, SD = 0.10; Elba2 mean = −0.76, SD = 0.11; Insv mean = −0.68, SD = 0.10 and CP190 mean = −0.63, SD = 0.11) (Figure 6B, cluster 2, Supplemental Table 7). Functional analysis of cluster 2 showed an overrepresentation of genes involved in developmental and morphological progresses (Figure 6D). As developmental and morphological associated genes were identified by Ueberschär et al. (2019) to be controlled by the Elba-complex, we compared our data with their ChIP-data for Elba1-3 and Insv (Figure 6G, S9A-F). Importantly, we found an overlap between our cluster 2 and genes associated to Elba1/3 ChIP-peaks. Together, our results strengthen the notion that the Elba complex is needed for distinct gene expression, and when downregulated by heat shock or mutation, there is a transcriptional de-repression at MBT.

**Figure 6.**
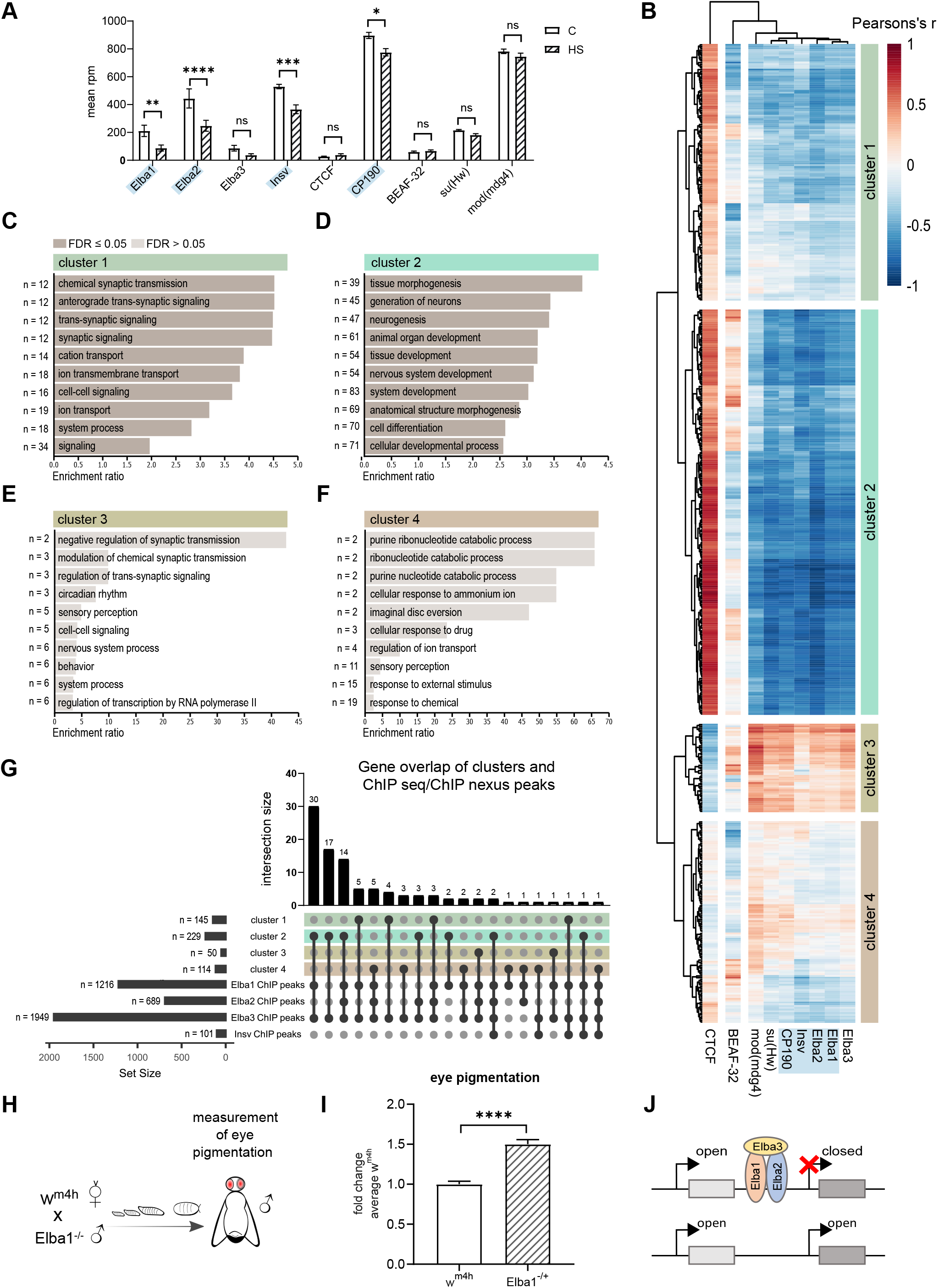
The Elba insulator binding complex is negatively associated with developmental genes and acts as a Su(var) for *w^m4h^*. **A.** Expression of known insulator binding factors in control and heat shocked embryos. The expression of Elba1, Elba2, Insv and CP190 decreases after heat shock exposure in the sensitive period. **B.** Correlations (Pearson’s r) between the factors described in (A) and all significantly upregulated genes with a fold change >1. Euclidean clustering shows that cluster 2 correlates inversely with most insulator binding factors, including the Elba family. Down-regulated factors are marked in blue. **C-F.** Gene ontology enrichment analysis of gene clusters from (B) was done using WebGestalt (Wang et al., 2017). Top 10 hits from over-representation analysis (ORA) for biological processes are presented per cluster. Genes from cluster 2 (C) are functionally enriched in developmental processes. **G.** Number of intersecting genes between the clusters identified in (B) and genes associated with previously published binding sites for Elba factors and Insv. 69 genes from cluster 2 (approx. 30% of cluster 2) overlapped with Elba factor binding sites, while much fewer genes (3) intersect with Insv binding sites. Cluster 1, 3 and 4 have almost no overlaps. Binding sites and their associated genes were acquired from published ChIP-seq/nexus data (Ueberschär et al., 2019). **H.** Virgin *w^m4h^* females were crossed with Elba1^-/-^ males, after which eye pigmentation was measured in 5 days old adult male offspring. **I.** Eye pigmentation in relation to average *w^m4h^*. Elba1^-/+^ had more pigmented eyes than control *w^m4h^*, indicative of a more open chromatin at this locus. Heads were measured in groups of 6-10 and normalized per fly. **J.** Schematic hypothesis of how the Elba insulator binding complex suppress position effect variegation in *w^m4h^* by partition of chromatin states and gene expression. 24 single embryos were analyzed per condition and presented with ± SEM in (A). * (p ≤ 0.05), ** (p ≤ 0.01), *** (p ≤ 0.001) and **** (p ≤ 0.0001) using multiple t-test with discovery by the two-stage linear set-up procedure of Benjamini, Krieger and Yekutieli. 47 controls and 50 Elba1 mutant samples were measured in (I), and presented with ± SEM, *** (p ≤ 0.001) using two-tailed t-test.

To test if reduced Elba1 during MBT would result in long-term epigenetic effects in the adult fly, we next crossed female virgins from the *w^m4h^* PEV-strain with homozygous *Elba1* mutant males (Figure 6H). Intriguingly, eye pigmentation of 5 days old males showed a dramatic loss of *white* silencing, thus mimicking the effect of an early embryonic heat shock (Figure 6I). In all, our study points to a central role for miRNA-Elba dependent fine-tuning of the emerging chromatin landscape, that—if disrupted—may have life-long consequences on the phenotype.

## DISCUSSION

Here, we provide new insights into the dynamics of sncRNA during the earliest stages of *Drosophila* embryogenesis as well as their response to heat shock. We found that heat shock induced an extensive increase of both maternal and zygotic miRNA, with the important exception of the mir-309 cluster—the master clock in temporal control of early embryogenesis. This early embryonic heat shock was echoed in the adult fly as reduced H3K9me3-dependent heterochromatin reported by the PEV-strain *w^m4h^*. Combining transcriptome-wide data of both sncRNA and long RNA from the same single embryos at the time of *de novo* chromatin state formation (stage 5) revealed strong associations between heat shock-induced upregulation of a specific group of miRNA (e.g. mir-2, mir-190 and bantam) and reduction of a specific gene cluster, where of 9 pre-MBT genes are only expressed at this point during *Drosophila* development (Chen et al., 2013). One of these genes, a newly described insulator binding factor—Elba1, acts as a transcriptional repressor to ensure correct gene expression during early development (Ueberschär et al., 2019). In line with this function, we find that a heat shock in the first hour of embryogenesis results in upregulation of genes involved in developmental patterning, and most interestingly, a substantial overlap between our upregulated genes and ChIP-data from Elba family of proteins. Finally, reduction of Elba1 efficiently mimicked the original effect on *w^m4h^* caused by the embryonic heat shock. Thus, our results suggest a miRNA-driven control of the zygotes first transcriptome to set the tone for forthcoming gene expression.

### Zygotic transcription of miRNA precedes the transcription of protein-coding genes

The first two hours of *Drosophila* embryogenesis is a dynamic period with extremely rapid mitotic cycles that are enabled by maternally loaded components. Simultaneous with the rapid expansion of genomic material, the zygote prepares to become self-sufficient by degrading maternal transcripts and organizing the chromatin architecture to enable stable endogenous transcription (Hamm and Harrison, 2018; Hug et al., 2017; Kwasnieski et al., 2019; Nien et al., 2011; Rudolph et al., 2007; Seller et al., 2019; Yuan and O’Farrell, 2016). Zygotic gene transcription is mainly initiated at embryonic stage 5, at the same time as heterochromatin is first detectable through microscopy (Rudolph et al., 2007), and the first tendency of chromatin organization in the form of topology associated domains (TAD’s) is evident (Hug et al., 2017). This is, however, proceeded by the nuclear appearance of the pioneering transcription factor Zelda (Nien et al., 2011), expression of non-paused pre-MBT genes (Chen et al., 2013), and massive amounts of aborted transcripts (Kwasnieski et al., 2019), suggesting that the transcriptional machinery is active but impeded by too rapid nuclear division (Edgar and Schubiger, 1986). We find the Zelda controlled miR-309 cluster to be significantly increased already between embryonic stage 1 and 2, long before the major initiation of zygotic gene transcription. At this time-point, the interphases are less than 4 min (reviewed in (Yuan et al., 2016)). Further, we see a dramatic increase of the miR-309 cluster as the cell cycles slow down to interphases of 8-14 min in embryonic stage 4 (Foe and Alberts, 1983), when preparing for gastrulation in stage 5.

It has been reported that the first genes to be transcribed are short. As an example, the mature Elba1 transcripts are not detected until nuclear cycle 13 (stage 4), whilst aborted Elba1 transcripts has been detected already at nuclear cycle 7-9 (stage 3) (Kwasnieski et al., 2019). As the short mitotic cycles still allows for these short transcripts, it would be likely that transcription of miRNA precursors of a few hundred nucleotides precedes the transcription of most mRNA, simply by the size difference.

### The making of a heterochromatin-based epigenetic memory

The position-effect-variegation strain *w^m4h^* has been extensively used for epigenetic research and enabled the discovery of multiple Su(var)s and E(var)s. The level of *w^m4h^* eye pigmentation, interpreted as gene-expression state at the *white* locus that depends on the heterochromatic strength, is set very early in development and kept during adulthood (Bughio et al., 2019). However, Bughio *et al*. showed that the variegation of the eye color in *w^m4h^* is not specific to the locus of *white*, but rather the consequence of a general heterochromatic landscape, where the silencing of *white* depends on the continuous maintenance of a nearby heterochromatic block. In line with previous research, we found that the most sensitive period to modulate this heterochromatic strength is before the MBT (Bughio et al., 2019; Hartmann-Goldstein, 1967; Lu et al., 1998; Seong et al., 2011). Thus, the setting of heterochromatic strength must be preceded by molecular events that are not depending on histone modification and 3D organization, but rather dictating it.

Our data suggest that environmental-sensitive miRNA that modulates the first zygotic transcripts coding for chromatin boundary elements and insulator binding factors could be such an initiating event. It could be argued that the Elba complex, with such extreme temporal restriction to mitotic cycle 13-14 (embryonic stage 4-5), could not have long-term epigenetic effects on the *white* gene, which is expressed first during late pupa stages. We find that Elba1, despite being expressed so transiently, has a suppressing effect on PEV in *w^m4h^*. Elba1 has been identified as an insulator binding factor as well as an repressor of transcription (Dai et al., 2015; Ueberschär et al., 2019). The Elba complex is tightly restricted to early development around MBT and declines at gastrulation (Figure S8) (Singer and Lengyel, 1997). During its active period, the Elba complex ensures gene expression integrity within the Bithorax complex (Aoki et al., 2012). Knock-down of the Elba complex does not only lead to loss of expression integrity of important developmental genes, but also to non-specific elevated gene expression (Ueberschär et al., 2019). Since we observed elevated gene expression as well as suppression of PEV variegation following heat shock, we hypothesize that heat shocked embryos may suffer from lower insulating capacity. Heat shock in *Drosophila* cell lines has been reported to weaken TAD border strength enabling relocation of architectural proteins, such as insulator binding elements, from TAD borders to inside TADs (Li et al., 2015). Heat shock during the sensitive pre-MBT period could therefore not only lead to reduced Elba1 (hence a less functional Elba complex) but also to less defined compartmentalization of chromatin. The consequences of a reduced expression of insulating binding factors, or their displacement, are potentially greater during this embryonic period than others as the initial TAD structures establishes at ZGA and are to a high extent maintained throughout development (Hug et al., 2017).

In conclusion, we have discovered that an early embryonic heat shock results in elevated levels of specific miRNA and reduced levels of zygotic insulator binding factors, including Elba1. Reduction of Elba1 correlates to upregulation of early developmental genes and promotes a more open chromatin structure in the adult fly. Our data fits a model where the initial heterochromatic tone is decided within the first two hours of embryogenesis via an extremely early miRNA-controlled insulator axis, where loss of insulation may deregulate early zygotic transcription and the transition to zygotic independence.

## ACKNOWLEDGMENTS

This research was supported by the by Swedish VR 2015-03141, Ragnar Söderberg’s foundation and Alice och Knut Wallenbergs foundation 2015.0165. The authors are grateful to Qi Dai’s lab for kindly sharing their Elba mutant fly strains. We would also like to acknowledge the staff at Linköping University core facility, especially Åsa Schippert and Anette Molbaek for their great support, and also Andrew S Belmont (University of Illinois) for his great linguistic support on the manuscript.

## AUTHOR CONTRIBUTIONS

Conceptualization, A.Ö, L.Ö and D.N.; Methodology, U.K, L.Ö and D.N; Investigation, L.Ö, S.S and H.E; Software, D.N, L.Ö and S.S; Validation, L.Ö; Visualization, L.Ö; Project administration, A.Ö, L.Ö; Writing – Original Draft, L.Ö, A.Ö and D.N; Funding Acquisition, A.Ö; Resources, A.Ö, M.R, R.R, H.E and U.K.; Data curation, L.Ö and D.N; Formal analysis, L.Ö and D.N; Supervision, A.Ö.

## SUPPLEMENTARY FIGURE LEGENDS

**Figure S1.**
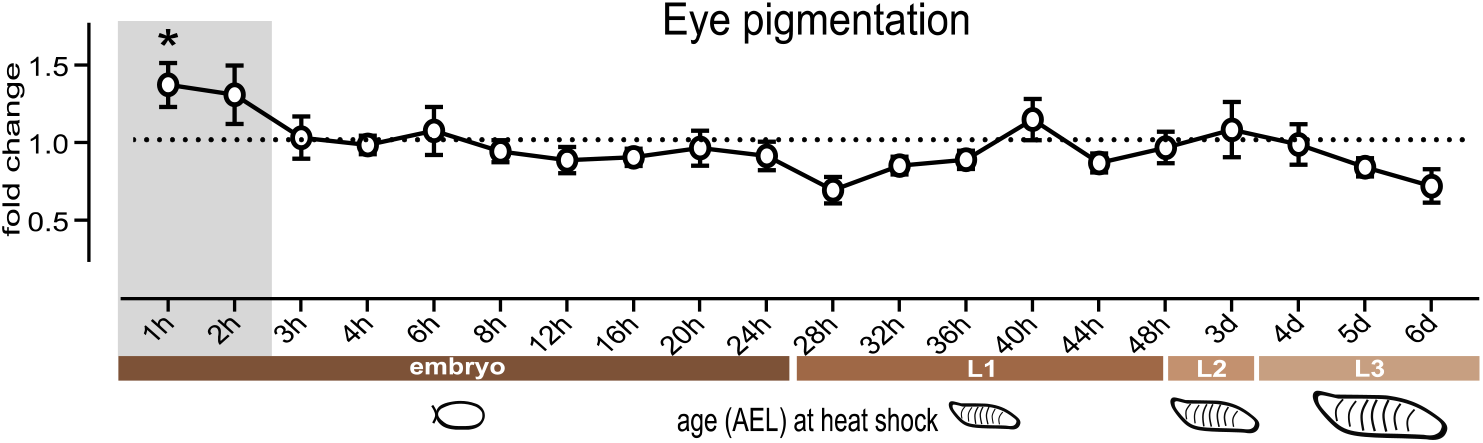
The most sensitive period for stress-induced epigenetic program is found in early embryogenesis. Spectrophotometric measurement of the heterochromatin eye pigment reporter *white* in 5 day old adult male *Drosophila w^m4h^* after one session of 1 h heat shock at 37° C performed during different embryonic and larvae stages of development. Graph represents eye pigmentation in relation to the average optical density of controls (not exposed to heat shock). We found the most sensitive period during early embryogenesis. Heads were measured in groups of 3-10 and normalized to the average optical density per head. n = 7. Presented with ± SEM, * (p ≤ 0.05) with ordinary one-way ANOVA with Dunnett’s multiple comparison test.

**Figure S2.**
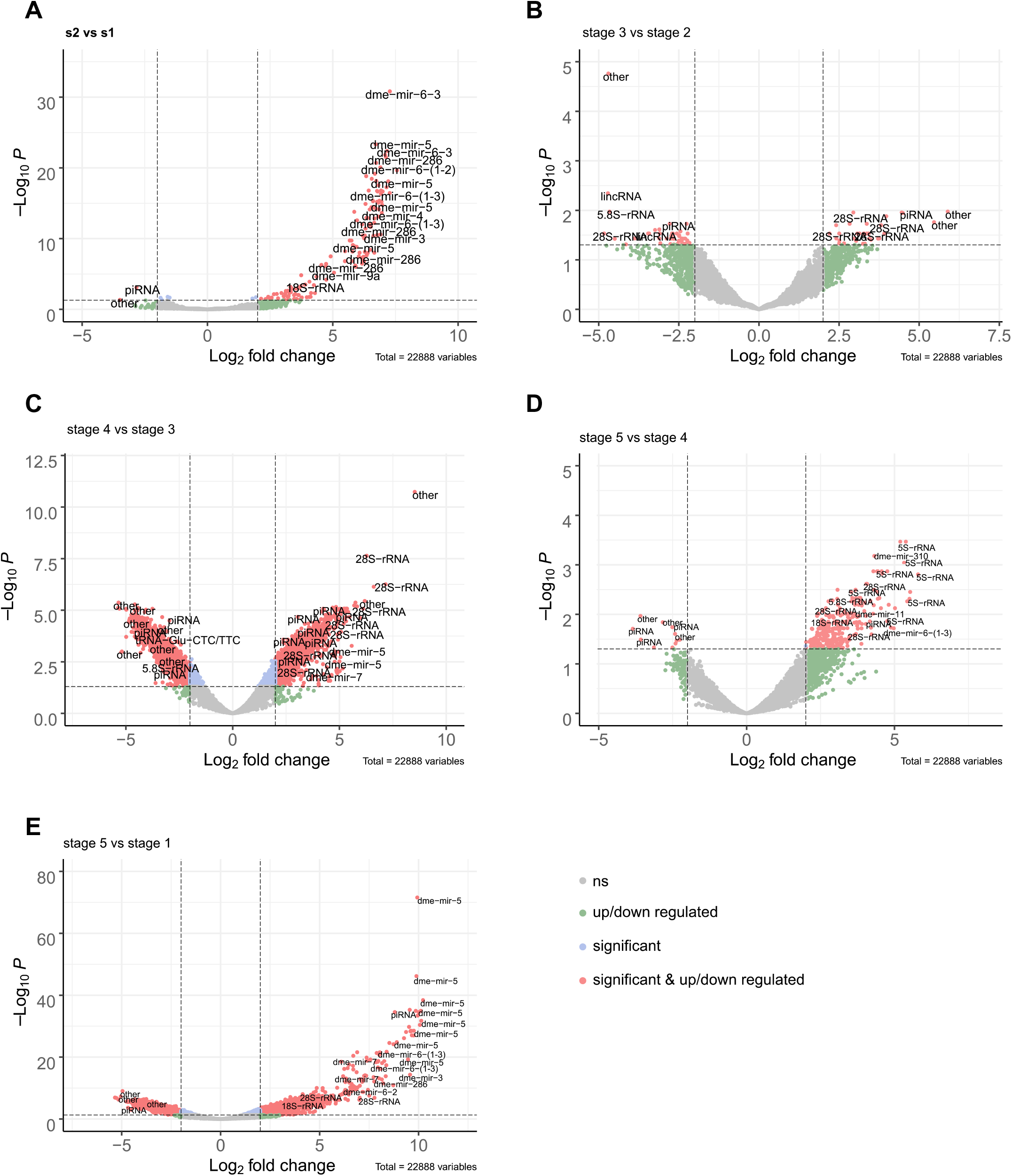
Volcano plots of differentially expressed sncRNA reads from stage to stage. Volcano plot showing differentially expressed unique sncRNA reads. Red indicates significance at p ≤ 0.05 (FDR corrected p-values) and a log2 fold change ≥ ± 1.5. **A.** Stage 2 vs stage 1 shows that there is an upregulation of the mir-309 cluster already between these stages. **B**. Stage 3 vs stage 2, **C.** Stage 4 vs stage 3. **D.** Stage 5 vs stage 4. **E.** Stage 5 vs stage 1. Groups of 4-5 single embryos were sequenced per stage.

**Figure S3.**
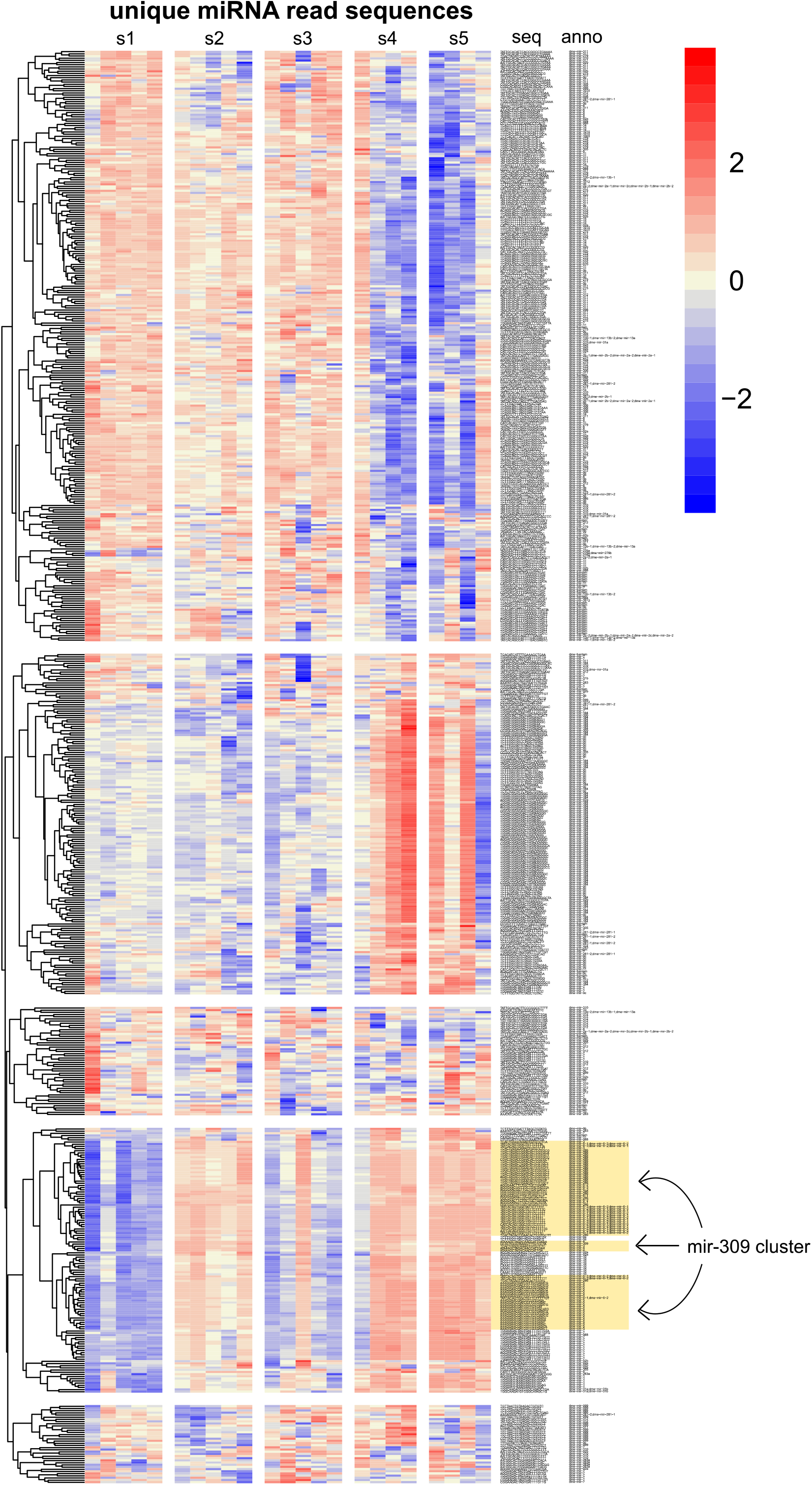
Heatmap over unique miRNA reads per embryo stage. Heatmap of all unique sequences aligning to miRNA per indicated stage. Columns represents unique samples (4-5 single embryos per stage). We detected a cluster of miRNA, including the mir-309 cluster (highlighted), that was increased between stage 1 and 2. Color bar represents the relative rpm expression per miRNA sequence.

**Figure S4.**
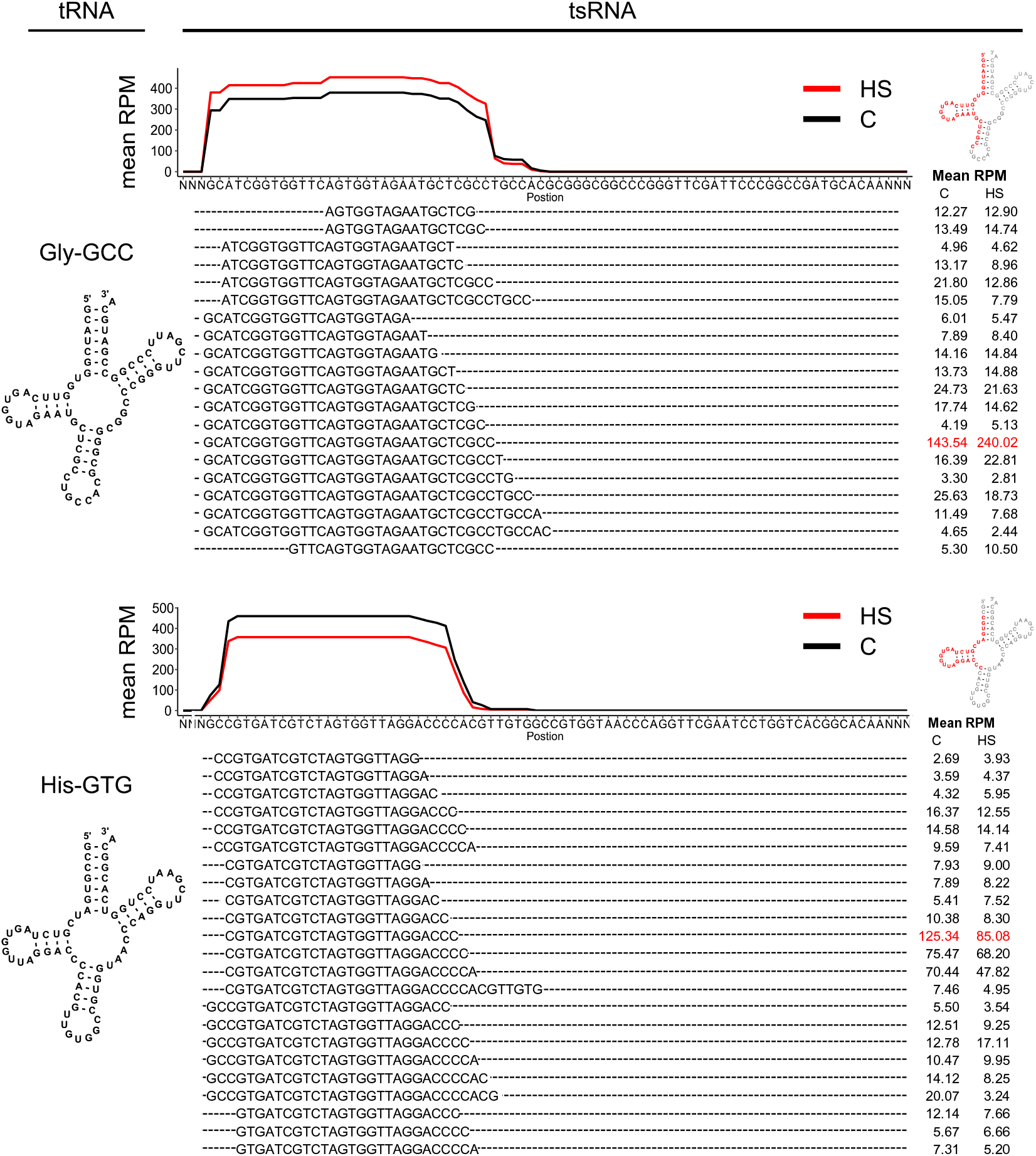
Heat shock leads to more tRNA-Gly-GCC halves and less tRNA-His-GTG fragments. Line graph shows mean rpm coverage of reads mapping to tRNA-Gly-GCC (upper) and tRNA-His-GTG (bottom). Mean RPM of each unique sequence contributing to the line plots are presented below each graph. The most differentially expressed sequences are marked in red.

**Figure S5.**
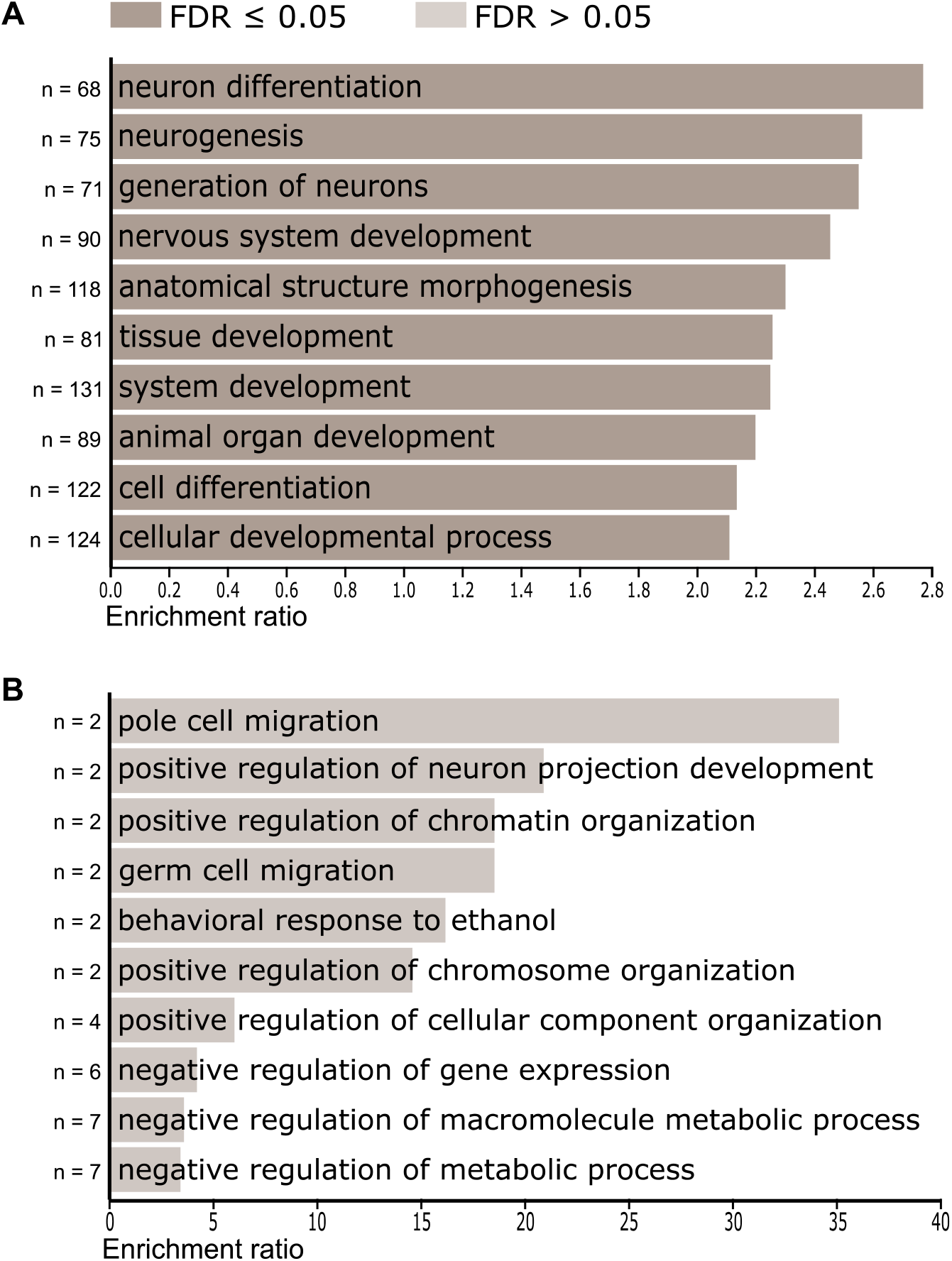
GO-term analysis of up and down regulated genes after heat shock. All significant genes (FDR corrected p ≤ 0.05) with log2 fold change ≥ 1 (A) or ≤ −1 (B) after exposure to heat shock during the sensitive period. **A.** Upregulated genes (n = 536) were enriched for developmental processes. **B.** Downregulated genes (n=42) showed no enrichment. Data analysed using WebGestalt (Wang et al., 2017). Top 10 hits from over-representation analysis (ORA) are presented per cluster.

**Figure S6.**
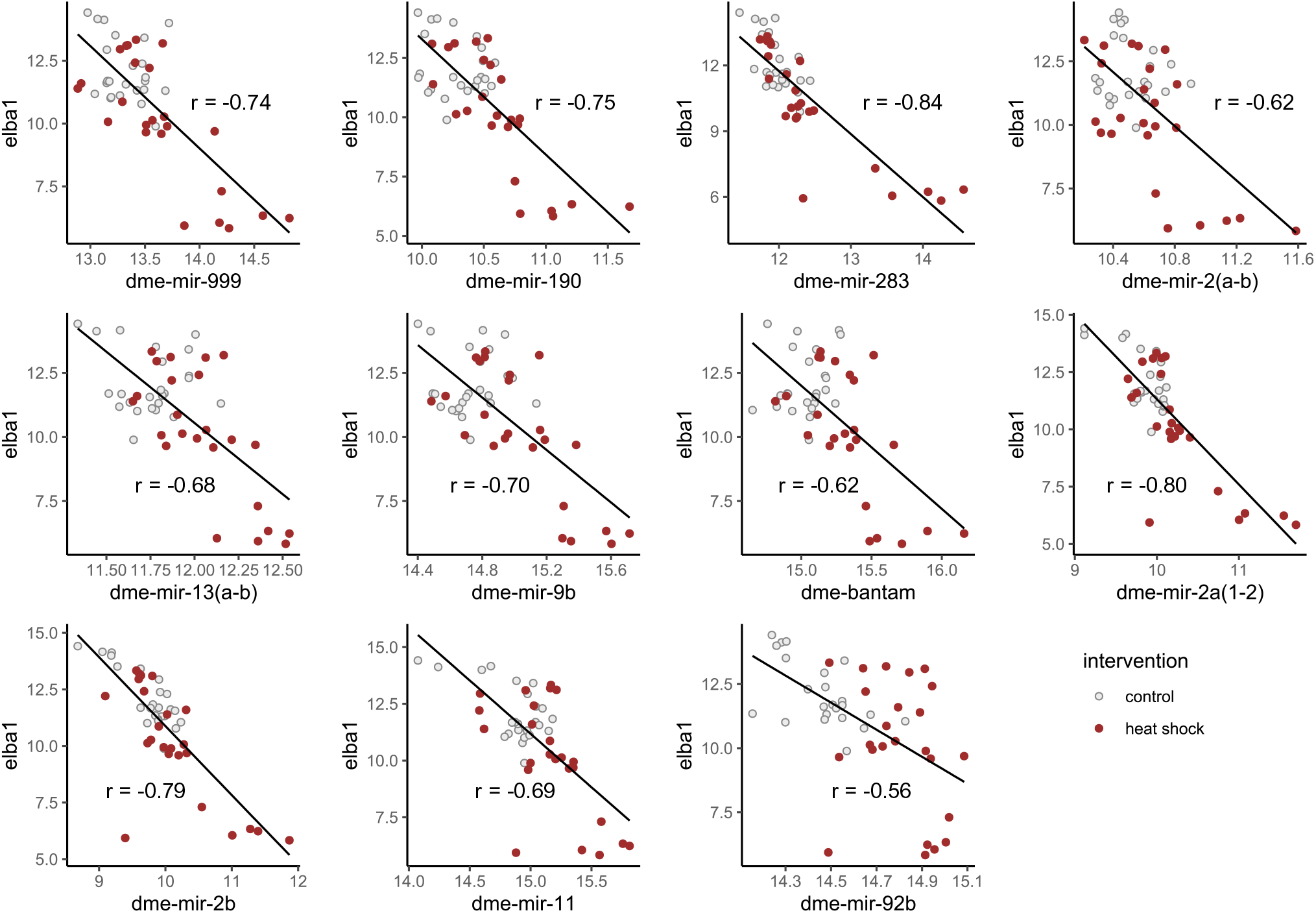
Strong negative correlation between upregulated miRNA and Elba1. Correlation between Elba1 and upregulated miRNA vst normalized counts. Red circles are heat shocked samples, grey circles are controls. Pearson’s r is associated with each graph. All p-values are < 0.0001.

**Figure S7.**
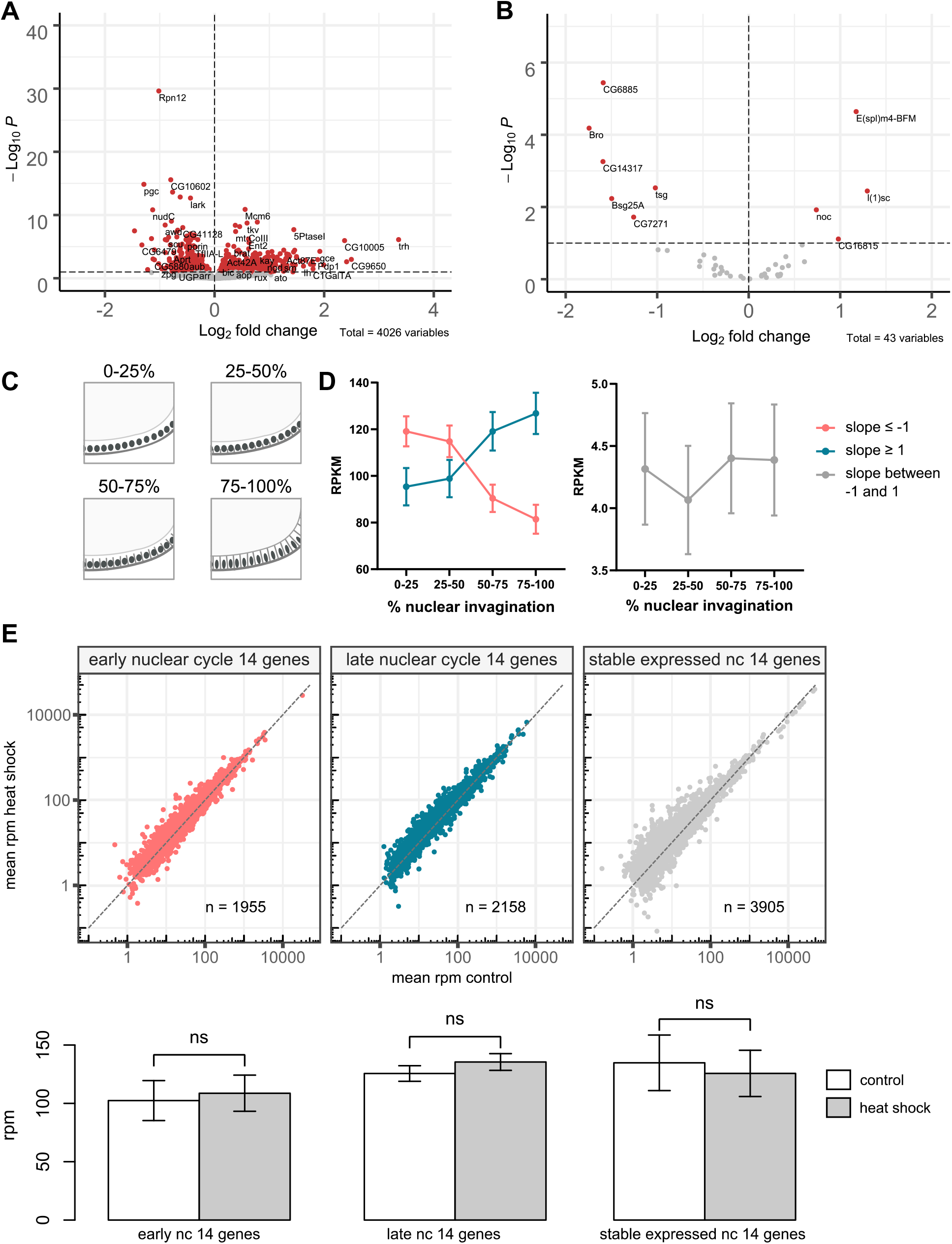
No sampling bias defected between heat shocked and control embryos. **A.** Maternally provided genes (classified in (Lott et al., 2011)) showed an equal distribution of up (n = 2133) respectively down (n = 1893) regulated genes. **B.** Early zygotic genes (1-2 h) (classified in (De Renzis et al., 2007)) showed an equal distribution of up (n = 21) respectively down (n = 22) regulated genes. **C-D.** Linear regression of genes expressed at 4 chronological divided parts of nuclear cycle (nc) 14, divided by percent of nuclear invagination (Lott et al., 2011), were used to define genes expressed early (slope ≥ 1), late (slope ≤ −1) or stable (slope between −1 and 1) during this cycle. **D.** Mean RPKM of genes per subdivided nc 14 classified as early, late or stable. Data from (Lott et al., 2011). **E.** Using this categorization, we compared gene expression between groups and found no significant changes (unpaired, two-tailed t-test).

**Figure S8.**
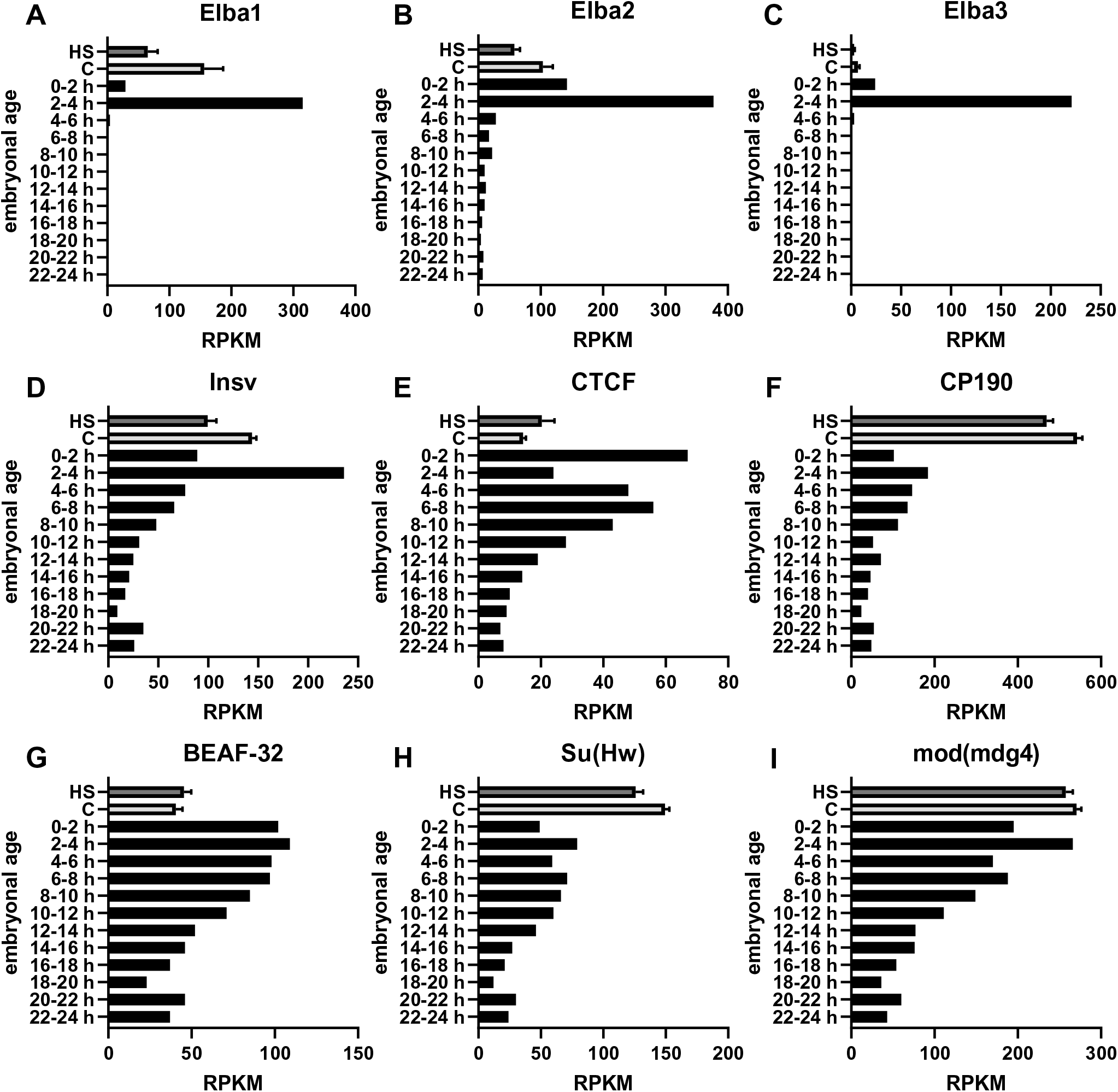
Embryonic expression of insulator binding factors. Expression levels of known insulator binding factors at different embryonic time-points. **A-C**. ELBA factors are specifically expressed (especially Elba1 and ELBA3) during pre-MBT and MBT. **D.** Insv is highly expressed in 2-4 h old embryos, but continuous to be expressed throughout embryogenesis. **E-I.** The other factors are also expressed throughout embryonic development but with a declining trend. Data was extracted from modENCODE.

**Figure S9.**
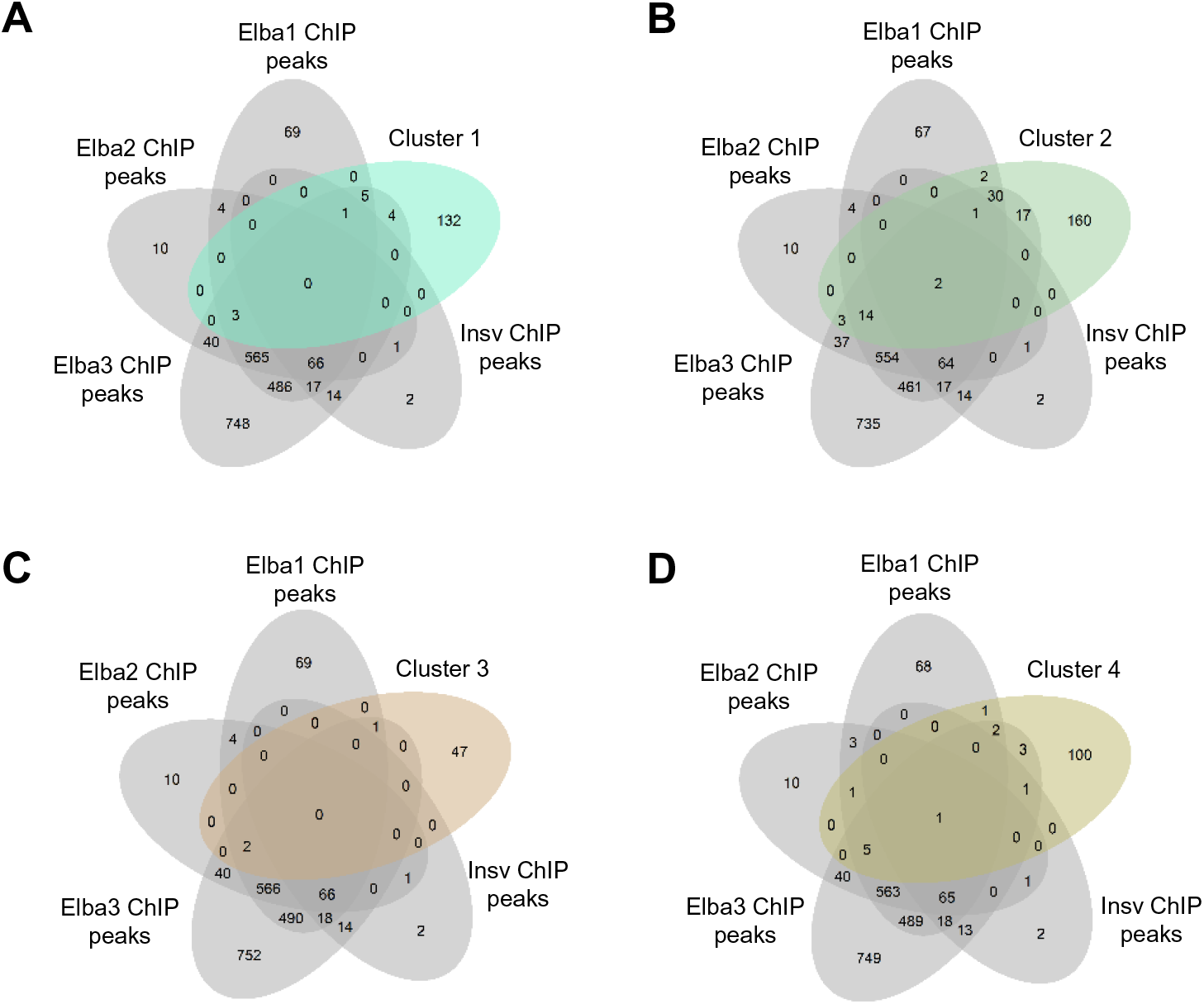
Occurrence of upregulated genes after heat shock in insulator binding factor binding-sites. Venn diagrams showing overlaps between gene clusters identified in figure 6B, and genes associated with published ChIP seq/ChIP nexus peaks of the Elba family and Insv insulator binding factors (Ueberschär et al., 2019). Cluster 2 shows more overlap with genes associated to binding sites for the Elba family than the other clusters.

### Supplementary data tables

**Supplemental_Table_S1:** Unique sequencing sample IDs for sncRNA and long RNA.

**Supplemental_Table_S2:** Reads per million (rpm) expression of unique sncRNA sequences obtained from stage 1-5 *Drosophila ln(1)w^m4h^* embryos.

**Supplemental_Table_S3:** Reads per million (rpm) normalized counts of all significant (FDR corrected p-value ≤ 0.05) unique sncRNA sequences in control and heat shocked *Drosophila ln(1)w^m4h^* embryos.

**Supplemental_Table_S4:** Reads per million (rpm) normalized sncRNA counts summarized per annotation feature of control and heat shocked *Drosophila ln(1)w^m4h^* embryos.

**Supplemental_Table_S5:** Reads per million (rpm) normalized significantly up-or downregulated genes of control and heat shocked *Drosophila ln(1)w^m4h^* embryos. Cluster numbers (figure 5D and 6B) are notated.

**Supplemental_Table_S6:** Correlation (Pearson’s r) for significantly changed miRNA and downregulated genes (belongs to figure 5).

**Supplemental_Table_S7:** Pearson’s r scores for significantly upregulated genes and insulator binding factors (belongs to figure 6).

**Supplemental_Table_S8:** Annotation sources for sncRNA.

## MATERIAL AND METHODS

### Fly husbandry

*ln(1)w^m4h^* (Gunter Reuter) *Drosophila* were maintained in a climate controlled 22°C incubator and kept on standardized food. Flies used for experiments were inbred for > 10 generations and flies with complete loss of PEV were not used for crossing. Elba1 mutant flies was kindly provided from Qi Dai’s lab (Ueberschär et al., 2019), and kept at room temperature (at approximately 22°C) on standardized food. Elba1^-/-^ - *ln(1)w^m4h^* crossings were kept in climate controlled 22°C incubator on standardized food.

### Eye pigment measurement

For screening of sensitive periods: eggs were collected on juice agar plates in tight intervals (30 min – 1 h) and exposed to one heat shock session at 37°C, or were kept as controls. For PEV expression in Elba1 mutants: virgin *ln(1)w^m4h^* were crossed with either Elba1 mutant males or *ln(1)w^m4h^* males and left to mate and lay eggs. Five or six different vials per crossing were set up and all were flipped 3 times. All experiments: flies were left to develop in climate controlled 22°C incubator. Males were decapitated 5 days after eclosure and their heads, collected in groups of 3-10, were first frozen in liquid nitrogen and then homogenized with a 5 mm ∅ metal bead (Qiagen) for 2 min at 40 Hz using TissueLyser LT (Qiagen). 500 µl PBS-tween (0.01%) was added and samples were shaken, kept in room temperature for 1 h and centrifuged. Absorbance at A480 was measured on supernatant using VersaMax (Molecular Devices) microplate reader. At least two biological replicates were collected per heat shock time and experiment, and heat shock experiments were performed 7 times.

### Sampling for sncRNA sequencing of developmental timeline

Eggs were collected on juice agar plates for 30 min and were immediately dechorionated. Staging was performed under SMZ 745 (Nikon) microscope using the criteria for Bownes’ stage 1-5 (Bownes, 1975). Single embryos were collected in 2 µl RNase free water with Recombinant RNase inhibitor (TAKARA) and ruptured with an RNase free needle. One 5 mm ∅ metal bead (Qiagen) and 500 µl Qiazol (Qiagen) was added per sample and shaken. n = 5 of stage 1-3 and n = 4 of stage 4 and 5.

### Sampling for sncRNA and long RNA sequencing after exposure to heat shock

Eggs were collected on juice agar plates in 30 min intervals and immediately exposed to one session of heat shock at 37°C for 30 min or kept as controls. Embryos were thereafter kept in climate controlled 22°C incubator for approximately 2 hours, dechorionated and staged under SMZ 745 (Nikon) microscope using the criteria for Bownes’ stage 5 (Bownes, 1975), including formed cells at egg surface and round pole cells at the posterior axis. Single embryos were collected in 2 µl RNase free water with RNase inhibitor and bursted with a RNase free needle. One 5 mm ∅ metal bead (Qiagen) and 500 µl Qiazol (Qiagen) was added per sample and shaken. We sampled 24 embryos per condition.

### RNA extraction and small RNA library preparation

RNA was extracted using miRNeasy Micro Kit (Qiagen) according to manufactures protocol. Quality was confirmed using Agilent RNA 6000 Nano kit (Agilent) on the 2100 Bioanalyzer Instrument (Agilent) prior to storage at −70 °C. NEBNext Small RNA Library Prep Set for Illumina (New England Biolabs) was used for library preparation according to manufacturer’s protocol with some changes. We downscaled all samples to half volume and added 2SrRNA block oligo (5’-TAC AAC CCT CAA CCA TAT GTA GTC CAA GCA-SpcC3 3’; 10 µM) (Wickersheim and Blumenstiel, 2013) to a final concentration of 2.5 µM together with SR-RT primer (from kit). Primers and adaptors from the kit were diluted 1:4 until PCR amplification, according to starting RNA concentration. PCR amplification was run for 15 cycles and NEBNext Index1-24 primers for Illumina were used (New England Biolabs).

Libraries were cleaned using Agencourt AMPure XP (Beckman Coulter) and run on pre-casted 6% polyacrylamide Novex TBE gel (Invitrogen). Bands of sizes 140-170 bp were selected. Gel extraction was made by centrifugation at 15 000 x g using gel breaker tubes (IST Engineering Inc) in DNA Gel Elution Buffer provided in the NEBNext kit. Samples were incubated at 37°C for 1 h on shaker, frozen in −70°C for 15 min and incubated in 37°C on shaker for 1 h once more. Gel debris was removed by Spin-X 0.45 µm tube. Libraries were precipitated overnight at −70°C in 1 µl Glycoblue (Invitrogen), 0.1 x volume of 3M Acetate (pH 5.5) and 3 x volume of 100% ethanol.Library sizes were measured on 2100 Bioanalyzer instrument (Agilent) using the Agilent High Sensitivity DNA kit (Agilent) and concentration was determined using QuantiFluor ONE ds DNAsystem on Quantus fluorometer (Promega). Equal concentrations of libraries were pooled and sequenced on NextSeq 500 sequencer using NextSeq 500/550 High Output Kit v2 with 75 cycles (Illumina). Unique sample IDs are summarized in Supplemental Table 1.

### Preprocessing of sncRNA sequencing results

We used Cutadapt version 1.18 to trim the adaptor sequence (AGATCGGAAGAGCACACGTCTGAACTCCAGTCACAT) from sncRNA reads and FastQC v.0.11.5 for quality filtering. Reads between 14- and 80 nucleotides, containing adaptor and with more than 80% of the bases having a phred quality score (Q-score) > 20 were retained. Mean sequence depth was 18.06 M reads (min = 12.01 M, max = 38.36 M) for data from the developmental timeline and 15.98 M reads (min = 12.15 M, max = 20.21 M) for heat shock experiments.

Trimmed reads were further mapped using SPORTS pipeline version 1.0.5 (Shi et al., 2018) with standard settings except following modifications; we replaced Rfam with repeatmasker (see supplementary methods), and included this at the bottom of the hierarchy. Within this pipeline, Bowtie version 1.1.2 was used returning one single alignment per read (with 0 mismatches if possible) with one mismatch allowed; -M 1 - -strata --best -v 1. Alignment was performed using the following hierarchy; reference genome, miRNA, tRNA, rRNA, piRNA, other ncRNA, repeats. For details and annotation sources, see supplementary table 8. Number of reads, length and annotation was retained per unique sequence.

### sncRNA seq analysis

A list containing experimental metadata, annotation information and count table was retained and filtered (min 20 reads in 17 % of samples for timeline experiment and 20 reads in 50 % of samples for heat shock experiment). An additional filter removing sequences with less than 0.01 rpm per sequence (in 17% of samples for timeline experiment or in 100 % of samples for heat shock experiment) was applied and information about sncRNA class was retained from annotation information.

### Long RNA library preparation

DNA was digested from aliquots from the same RNA extracted as described above (RNA extraction and smallRNA library preparation) with RNase-Free DNase Set (Qiagen) according to kit protocol and concentrated using Oligo Clean & Concentrator (Zymo research) according to kit protocol but adjusted for sample volumes. RNA quality was determined on the 2100 Bioanalyzer Instrument (Agilent) using RNA 6000 Nano kit (Agilent).

cDNA was synthesized using Ovation RNA-Seq Systems 1-16 for model organisms (NuGEN) according to kit protocol. Samples were sonicated 6 times in 15 sec on- 15 sec off intervals using Bioruptor Pico sonication device (diagenode). Library construction was done using the Ovation RNA-Seq Systems 1-16 for model organisms (NuGEN) according to protocol, and cycles for library amplification was determined using 7900HT Fast Real-Time PCR System (Applied Biosystems™). The amplification buffer and enzyme mixes provided in library kit was used for the master mix together with EvaGreen for qPCR (Biotium).

Libraries were amplified according to mean cycle for exponential PCR amplification per experiment (16 cycles), and purified according to protocol. Library sizes were measured on the 2100 Bioanalyzer Instrument (Agilent) using High Sensitivity DNA chip (Agilent) and concentration was determined using QuantiFluor ONE ds DNAsystem on Quantus fluorometer (Promega). Equal concentrations of libraries were pooled and sequenced on NextSeq 500 sequencer using NextSeq 500/550 High Output Kit v2 with 75 cycles (Illumina). Unique sample IDs are summarized in Supplemental Table 1.

### Preprocessing and analysis of long RNA sequencing results

We used Cutadapt version 1.18 to trim the adaptor sequence (AGATCGGAAGAGCACACGTC) from long RNA reads and FastQC v.0.11.5 for quality filtering. Reads over 14 nucleotides and with more than 80 % of the bases having a phred quality score (Q-score) > 20 were retained. Depth per library was 21.54 M reads (min = 20.26 M, max = 31.66 M reads). STAR genome index files were generated using Drosophila_melanogaster.BDGP6.28.dna.toplevel fasta and Drosophila_melanogaster.BDGP6.28.101.gtf (Ensembl). These genome index files were then used on trimmed reads using STAR (v.2.5.0a) with standard settings and indexed using samtools (v.1.3.1). Standard featurecounts (v.1.5.0-p1) settings was used for assigning reads to genomic features, with a minimal overlap of 15 bases. A list containing experimental metadata, annotation information and countable was retained and filtered (min 10 reads in 50 % of samples).

In order to compare maternally loaded and early zygotic genes between control and heat shocked embryos, we extracted the genes in our data that matched the classifications made by Lott *et al*. (Dataset S1 in (Lott et al., 2011)) (maternal) or by De Renzis *et al*. (Table S8 in (De Renzis et al., 2007)) (early zygotic). To generate a matrix of correlation, we used the rcorr function with the Pearson’s option within the Hmisc package (version 4.2-0). We further used Euclidean clustering within the pheatmap package (version 1.0.12). For analysis of overlaps with pre-MBT classifications, we used classifications made by Chen *et al*. (Table 1 in (Chen et al., 2013)), to compare against genes from indicated gene clusters. For overlaps between heat shock induced upregulation of genes and Insv- and Elba factor binding sites, genes notated with corresponding binding sites in Ueberschär *et al*. (supplementary data 4 in (Ueberschär et al., 2019)) was extracted and compared to indicated clusters.

### Statistics

All statistical analysis was done in R 3.6.0, R 4.1.0 or GraphPad Prism v.8.4.3. For eye pigment statistical analysis, ordinary one-way ANOVA with Dunnett’s multiple comparison or two-tailed t-test was used as indicated. 1 outlier (data set used in 1C, at 12 h) was removed using the ROUT method (Q = 0.1%). As indicated, we used either rpm (Supplemental Table 2-5) or variance stabilizing transformation (vst) from DEseq2 (version 1.24.00) for normalization of sncRNA and long RNA sequencing results. For statistical analysis, DEseq2’s build in Wald test after negative binominal fitting, unpaired one- or two-tailed t-test, or multiple t-test (with discovery determined using the two-stage linear set-up procedure of Benjamini, Krieger and Yekutieli) was used.

### Data availability

The data sets generated during this study are available upon request.

